# Structural constraints and drivers of molecular evolution in a macromolecular complex; the kinetochore

**DOI:** 10.1101/2024.07.10.602950

**Authors:** Hannah K. Pare, Alexandra L. Nguyen, M. Sabrina Pankey, Iain M. Cheeseman, David C. Plachetzki

## Abstract

Evolutionary theory suggests that critical cellular structures should be subject to strong purifying selection as protein changes would result in inviability. However, how this evolutionary principle relates to multi-subunit complexes remains incompletely explored. For example, the macromolecular kinetochore complex, which mediates the faithful segregation of DNA during cell division, violates the expectation of purifying selection as subsets of kinetochore proteins exhibit rapid evolution despite its critical role. Here, we developed a multi-level approach to investigate the evolutionary dynamics of the kinetochore as a model for understanding how an essential multi-protein structure can experience high rates of diversifying selection while maintaining function. Our comprehensive approach analyzed 57 kinetochore genes for signatures of purifying and diversifying selection across 70 mammalian species. Intraspecies comparisons of kinetochore gene evolution showed that members of the order Afrotheria experience higher rates of diversifying selection than other mammalian orders. Among individual loci, genes that serve regulatory functions, such as the mitotic checkpoint genes, are conserved under strong purifying selection. In contrast, the proteins that serve as the structural base of the kinetochore, including the inner and outer kinetochore, evolve rapidly across species. We also demonstrated that diversifying selection is targeted to protein regions that lack clear structural predictions. Finally, we identified sites that exhibit corresponding trends in evolution across different genes, potentially providing evidence of compensatory evolution in this complex. Together, our study of the kinetochore reveals a potential avenue by which selection can alter the genes that comprise an essential cellular complex without compromising its function.

## INTRODUCTION

When comparing genomes, important functional elements are often defined as those with significant sequence conservation across evolutionary time, applying the logic that purifying selection acts against sequence changes that would disrupt function. The principles of sequence evolution become more complicated when considering macromolecular complexes that contain multiple protein subunits that interact with each other. Sequence evolution in one subunit may require compensatory evolution in a second subunit to maintain the function of the larger complex (1). In this case, sequence evolution in macromolecular subunit loci, if not highly constrained by purifying selection, could lead to a runaway evolutionary process that might ultimately destroy the function of the complex, or force macromolecular complexes to evolve so rapidly as to be incomparable across taxa. The fact that many such complexes are known to be conserved throughout macroevolution (kinetochore, spliceosome, ribosome etc.; (2–4) suggests the presence of mitigating factors that intercede in such runaway processes. Such factors could include variation in selection intensity across proteins that form the complex, where some subunits or functional modules, are permitted to explore sequence space whereas others are held tightly in check by purifying selection (5). In addition, variation in the types of protein structure where sequence change is more, or less, permissible could also allow sequence diversification and compensatory evolution to take place in some protein modules without initiating a runaway process (6). However, few studies have taken a comprehensive approach to studying the molecular evolution of macromolecular complexes, and such possibilities remain unclear.

One such complex is the kinetochore, which facilitates chromosome segregation to daughter cells during mitosis and meiosis. The kinetochore is subdivided into functional modules based on their localization within the complex and their function. The Constitutive Centromere-Associated Network (CCAN) resides at the base of the kinetochore and is comprised of 16 proteins that assemble on centromeric DNA constitutively to serve as the foundation for kinetochore assembly (7–9). Unlike many kinetochore proteins, the CCAN remains localized to centromeric DNA throughout the cell cycle. The outer kinetochore assembles upon the platform of the CCAN at mitotic entry to facilitate the attachment of the kinetochore to spindle microtubules. In mammalian cells, two arms of the CCAN comprised of CENP-C and CENP-T interact with outer kinetochore proteins to facilitate recruitment of the microtubule-binding NDC80 complex (10–14). Additional regulatory modules also associate with the kinetochore in mitosis, including the Chromosomal Passenger Complex (CPC), which regulates chromosome alignment and segregation (15), and the Mitotic Checkpoint Complex (MCC), which monitors kinetochore-microtubule attachments (16–18).

Most kinetochore genes show little pleiotropy, so selective pressure amongst individual loci can be attributed to a shared function in cell division. The critical role of the kinetochore in cell division, combined with the requisite cooperation of its more than 100 protein components suggests that changes over the course of evolution would be highly constrained. However, kinetochore proteins evolve at faster than expected rates, with human-mouse sequence identity as low as 39% for some orthologs. In contrast, some ribosomal proteins share greater than 80% sequence identity even between distantly related species such as zebrafish and humans (19). Therefore, the kinetochore represents a paradox: how does a macromolecular complex, encoded by > 100 loci, experience high rates of diversifying selection while maintaining its essential function?

Here, we used the kinetochore’s rapid evolutionary dynamics as a model to investigate how macromolecular complex sequences evolve. We analyzed a comprehensive set of 57 kinetochore genes across 70 mammalian species for signatures of purifying and diversifying selection. We show that kinetochore genes in Afrotheria, a clade of mammals originating in Africa including the elephant and the cape elephant shrew, experience significantly higher rates of selective pressure than other mammalian orders, specifically within CCAN and Outer Kinetochore modules. Our analysis recapitulates previous findings that CCAN genes experience strong diversifying selection together with the outer kinetochore genes (20). We further show that diversifying selection is enriched in unstructured regions across kinetochore proteins highlighting a possible route by which selection may alter protein sequence and/or function without perturbing other secondary structures required for essential protein-protein interactions. Finally, we identify sets of genes that have experienced independent episodes of diversifying selection at the same residues in multiple species. Such inter-protein corresponding residues occur in protein pairs that share functional connections, including the two functional arms that direct outer kinetochore assembly - CENP-T and CENP-C. We propose an evolutionary model and a partial resolution to the kinetochore paradox by which diversifying selection in a subset of proteins is compensated for by additional diversifying selection in unstructured regions of other proteins in the complex.

## RESULTS

### A comprehensive mammalian kinetochore dataset

To assess kinetochore evolution, we first created a dataset of genomic sequences across a diverse group of species. Given the substantial differences in kinetochore protein sequences across eukaryotes, we restricted our analysis to mammalian species, which include a range of species with high quality genomes. We included genome sequences for a dataset of 70 mammalian species and four outgroups (Figure 1B; Table S1) in an orthology analysis and identified groups of orthologs representing 57 mostly non-pleiotropic genes that represent the full scope of kinetochore structure and function. This included 15 genes that make up the CCAN, 13 genes that form the outer kinetochore, 13 checkpoint genes, the four genes that make up the CPC, and 12 genes that are not easily categorized in a specific functional group (referred to as miscellaneous) (Figure 1A).

**Figure 1.**
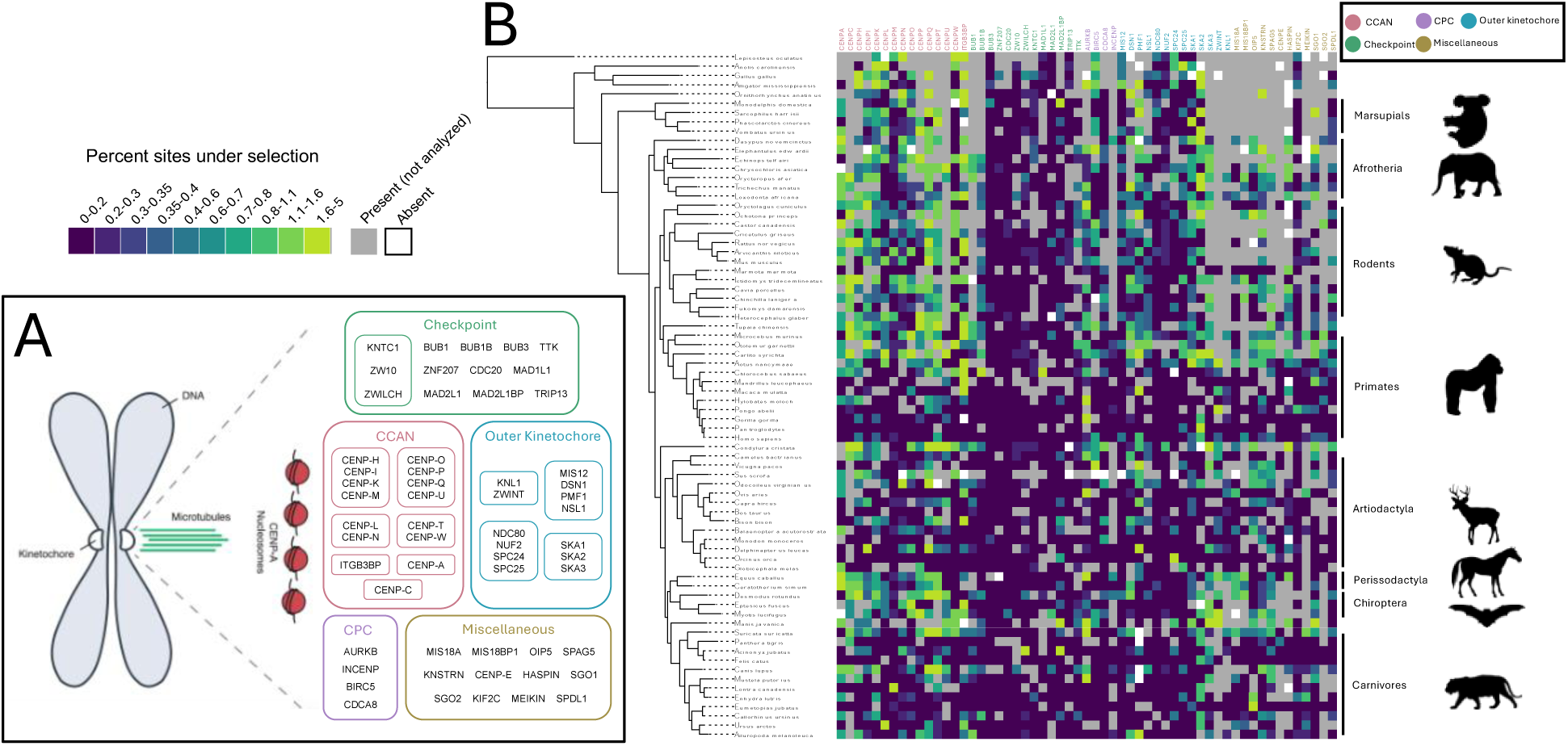
(A) 57 kinetochore genes were analyzed as a part of this study representing five functional groups (Checkpoint, CCAN, Outer kinetochore, CPC and Miscellaneous). Proteins that function in complex are circled. (B) The percentage of sites under episodic diversifying selection according to MEME for each gene (column) in each species (row) shown as a heatmap. Genes are organized by the functional groups presented in (A). Notable orders are highlighted. Genes absent from a particular species are represented by a white box; gray boxes represent genes that are identical in sequence to an orthologous gene of another species.

### The evolutionary dynamics of kinetochore proteins reveal clade-specific differences

We first sought to define the degree of selection in kinetochore genes among mammalian clades using the branch site model implemented in aBSREL (21). To assign evidence of selection to species lineages, we only considered selection at terminal branches. We tested for differences in the number of genes under episodic diversifying selection between orders with at least three representative species in our analysis (Carnivora, Chiroptera, Rodentia, Artiodactyla, Primates and Afrotheria) (Figure 1B). Using this model (aBSREL), the number of kinetochore genes under episodic diversifying selection was not significantly different between mammalian orders (ANOVA, F_5,51_ = 0.91, *p =* 0.482) (Figure S1). This was true even when testing for differences amongst genes within each functional group (*p* values between 0.348 and 0.743; see Figure 1A for functional categories). However, some species deviated from this trend, exhibiting an elevated number of genes under diversifying selection. For example, the Tasmanian devil (*Sarcophilus harrisii*) had the highest number of genes under selection (8 genes), followed by the little brown bat (*Myotis lucifugus*) (6 genes under selection) (Figure S1). Although aBSREL did detect selection on internal branches, most branches detected under this model were terminal branches (191 out of 247 total branches identified). Even when considering the total number of terminal versus internal branches, terminal branches were enriched for diversifying selection under this model (*p=* 2.2e-16, odds *=* 3.309) suggesting that most of the diversifying selection in kinetochore proteins occurred after the diversification of the mammalian orders.

It is possible that the low statistical power of our aBSREL-based approach could explain our inability to detect clade-specific differences. Therefore, we also used MEME (22), which tests for evidence of diversifying selection on each branch and for each site within the alignment, allowing for greater sensitivity across small sample sizes. We tested these data as above to search for clade-specific differences in the evolutionary dynamics of mammalian kinetochore loci. We found that the CCAN genes of afrotherians have a significantly higher number of sites under selection (lsmean = 3.31 sites per gene, CI: 2.69 – 3.93) when compared to primates (lsmean = 2.15, CI: 1.69 – 2.62) and carnivores (lsmean = 2.00, CI: 1.51 – 2.48) (ANOVA F_5, 506_ = 2.964, *p* = 0.012) (Figure 1B). Similarly, the outer kinetochore genes of afrotherians show significantly greater selection (lsmean = 2.30 sites under selection, CI: 1.898 - 2.71) when compared to the carnivores (lsmean = 1.32, CI: 0.989 - 1.66) (ANOVA F_5, 399_ = 6.74, *p* = 4.79e-06) (Figure 1B). We found no significant differences in the mean number of sites under selection across orders when looking at the checkpoint genes (ANOVA F_5, 391_ = 1.756, *p* = 0.121) and the CPC genes (ANOVA F_5, 117_ = 0.259, *p* = 0.935) (Figure 1B). When testing for differences within the miscellaneous category, there was a significant statistical interaction between the gene and the mammalian order variables (F_46, 256_ = 2.969, *p* = 2.67e-08). To eliminate the variation due to this statistical interaction, differences amongst orders were tested for each gene in this category. Of the 12 genes tested, two had significant differences in the number of sites under selection between orders - HASPIN (F_4,20_ = 4.188, *p* = 0.0127) and MIS18BP1 (F_4,8_ = 13.28, *p* = 0.00131). Afrotheria has more sites under selection (8.2 sites, CI: 5.56 – 10.84) in HASPIN than Artiodactyla (2.17, CI: -0.24 – 4.57) (Figure 1B). Chiroptera (16.0, CI: 10.34 – 21.66), Rodentia (13.5, CI: 9.50 – 17.50) and Afrotheria (10.0, CI: 6.00 – 14.00) have more sites under selection in MIS18BP1 than Carnivora (2.6, CI: 0.43 – 4.71) (Figure 1B).

Additionally, a few species displayed an exceptionally high numbers of sites under selection in specific genes, but these instances do not reflect clade-wide trends. For instance, when accounting for gene length, the white-tailed deer (*Odocoileus virginianus)* has experienced a notably high amount of selection in CENPU (21 sites), whereas the gorilla (*Gorilla gorilla*) and the Koala (*Phascolarctos cinereus*) possess high numbers of sites under selection in ITGB3BP (9 sites under selection in each species) (Figure 1B, S1). When combined across all kinetochore genes, the star-nosed mole (*Condylura cristata*) had the highest number of sites under selection (156 sites) and appears to be undergoing rapid diversifying selection across the kinetochore landscape (Figure 1B). MEME analyses also detected selection at internal branches, although most selection occurred in terminal branches. Together, analyses of clade-specific differences in diversifying selection among kinetochore genes suggest elevated diversifying selection in afrotherians compared to other mammalian orders, specifically within CCAN and outer kinetochore genes.

### Diversifying selection drives sequence changes in CCAN and outer kinetochore proteins

Next, we examined the selection in specific genes across mammalian species. We assessed each of the 57 kinetochore genes for episodic and pervasive diversifying selection using MEME (22), which identifies sites where a proportion of branches are evolving under diversifying selection (episodic diversifying selection), and FUBAR (23), which identifies sites under selection averaged across the alignment (pervasive diversifying selection). Estimates of episodic diversifying selection are more sensitive than pervasive diversifying selection approaches and, accordingly, we detected notably fewer sites under pervasive diversifying compared to episodic diversifying selection across all genes in our analysis (Figure S2; S3). In total, we identified 1,770 sites with evidence of episodic diversifying selection and only 104 sites with evidence of pervasive diversifying selection across the 32,042 sites considered amongst 57 kinetochore genes. Notably, all but six of the 108 sites identified by FUBAR as displaying pervasive diversifying selection were also identified by MEME as being under diversifying selection, further exemplifying the enhanced sensitivity of site-specific analyses. All genes in our analysis had at least one site under episodic diversifying selection (range: 1 - 102 sites, mean = 31, median= 23) (Figure S4A), and 43 of the 57 genes had at least one site under pervasive diversifying selection (range: 0 - 15 sites, mean = 1.82, median = 1) (Figure S4B). As expected based on the relative numbers of sites available, we identified a significant linear relationship between the length of a gene and the number of sites under episodic diversifying selection (*p* = 5.164e-10, R^2^ = 0.5074) amongst the 57 kinetochore loci (Figure S5).

We also analyzed the relationship between evolutionary rate and function. For these analyses, we considered the percentage of sites under episodic diversifying selection (number of episodic diversifying sites/total number of sites). We observed significant differences in the degree of episodic diversifying selection between functional groups (ANOVA *p* = 6.36e-05, F(_4,52_) = 7.646). The CCAN (lsmean = 10.78% sites under episodic diversifying selection, CL: 8.68 – 12.87%) and outer kinetochore genes (lsmean = 8.20%, CL: 5.95 - 10.45%) exhibit a higher mean percentage of sites under episodic diversifying selection when compared to the checkpoint genes (lsmean = 2.82%, CL: 0.57 - 5.07%) (Figure 2; S3). CCAN genes also have a significantly higher percentage of sites under episodic diversifying selection than the genes categorized as miscellaneous (lsmean = 5.25%, CL: 2.90 – 7.59%) (Figure 2; S3). Two outer kinetochore genes showed the highest percentage of sites under episodic diversifying selection: PMF1 (19.51%) and SKA2 (20.66%) (Figure 2; S4A; S6). They are followed by ITGB3BP (18.64%), CENPQ (16.04%), AURKB (15.07%) and CENPU (14.59%). Conversely, the 6 genes with the lowest percentage of sites under episodic diversifying selection each function as regulatory players in the Spindle Assembly Checkpoint signaling pathway (TRIP13: 0.23%, ZNF207: 0.43%, CDC20: 0.80%, BUB3: 0.92%, MAD2L1: 1.46% and ZW10: 2.05%) (Figure S4A).

**Figure 2.**
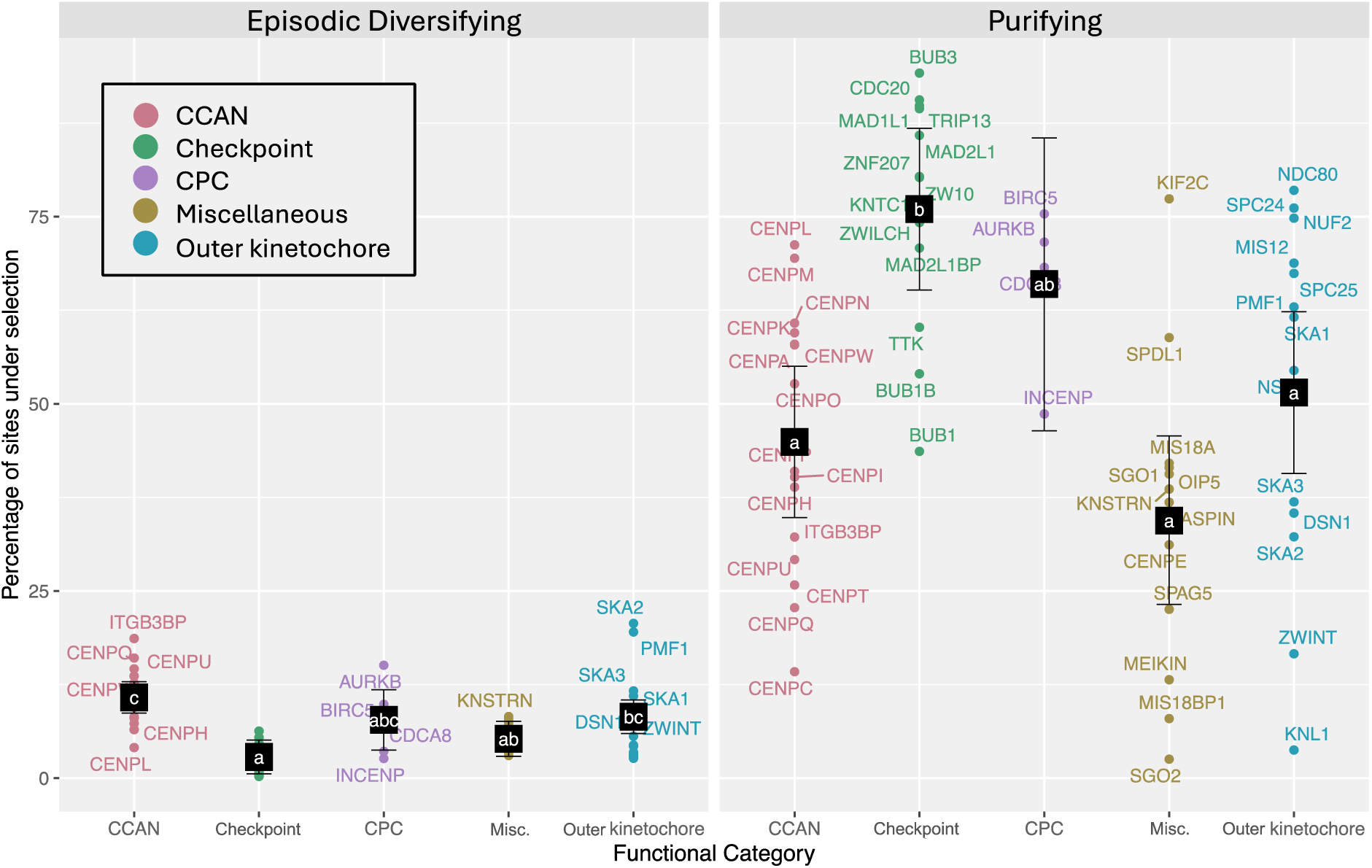
The percentage of sites under selection in each gene according to functional group (red: CCAN genes, green: checkpoint, purple: CPC, gold: miscellaneous and blue: outer kinetochore). Black squares are the least squares mean for each functional category and bars are the 95% confidence interval around the means. Letters represent significance groups according to a pairwise Tukey test comparing the means of all five groups.

CCAN genes also show a significantly higher mean percentage of sites under pervasive diversifying selection (lsmean = 0.988%, CL: 0.718 – 1.259%) than the checkpoint genes (lsmean = 0.134%, CL: -0.157 – 0.424%), miscellaneous genes (lsmean = 0.313%, CL: 0.010 – 0.615%), and outer kinetochore genes (lsmean = 0.412%, CL: 0.121 – 0.702%) (ANOVA *p* = 0.000423, F(_4,52_) = 6.096) (Figure S2; S3). The three genes with the highest percentage of sites under pervasive diversifying selection are CENPH (2.429%), CENPW (2.273%), and CENPI (1.984%), each of which are CCAN genes (Figure S4B; S6). They are followed by SKA3 (1.699%) and CENPM (1.667%). Notably, most proteins in the CENP-HIKM complex possess an above average percentage of sites under pervasive diversifying selection (H: 2.429%, I: 1.984%, K: 1.115%, M: 1.667%) (Figure S3; S4B; S6). Only CENP- K has an average that falls below the upper 95% confidence limit for the mean of CCAN genes (1.259%). Of the 57 genes in our analysis, 14 genes did not have any sites under pervasive diversifying selection (Figure S4B). In summary, we identified trends in the degree of diversifying selection according to the functional contribution of a gene within the kinetochore complex; genes that comprise the CCAN and outer kinetochore experience greater diversifying selection than checkpoint genes, which have very few sites under diversifying selection.

### Checkpoint proteins exhibit strong signals of purifying selection

Although our analyses of diversifying selection highlighted genes with the least amount of selectively evolving sites, a lack of diversifying selection does not itself equate to sequence conservation. To assess which genes are highly selectively conserved, we determined the number of sites under purifying selection using the program FUBAR (23). All 57 genes in our analysis exhibited sites under purifying selection (range 32 – 1,655 sites, mean = 260, median = 157) (Figure 2; S4C). Similar to our analysis of episodic diversifying selection, we found a significant linear relationship between gene length and the number of sites under purifying selection (*p* = 3.802e-07, R^2^ = 0.377) (Figure S7). As in our previous analysis, we accounted for this relationship by comparing the percentage of sites under purifying selection across genes. We found that the mean percentage of sites under purifying selection among checkpoint genes (76.0%, CL: 65.2 - 86.8%) is significantly higher than the mean for outer kinetochore (51.5%, CL: 40.7 – 62.3%), CCAN (44.9%, CL: 34.8 - 55.0%), and miscellaneous genes (34.4%, CL: 23.2 – 45.7%) (Figure 2; S3). As expected, many of the genes with very few sites under diversifying selection show the highest degree of purifying selection (Figure S4). The checkpoint components BUB3, CDC20, TRIP13, and ZNF207 are among the most conserved genes (BUB3: 94.17% of sites under purifying selection, CDC20: 90.58%, TRIP13: 89.81%, ZNF207: 80.35%) (Figure 2; S4C). Additionally, MAD1L1 and MAD2L1 both have a considerably high percentage of sites under purifying selection (89.42% and 85.85% respectively) (Figure 2; S4C). Conversely, SGO2 had the lowest percentage of sites under purifying selection (2.53%) (Figure 2; S4C). Altogether, the analyses of purifying selection highlight its role in maintaining sequence conservation of these genes, complementing our findings above that checkpoint genes exhibit few sites under diversifying selection.

### Intrinsically unstructured regions are enriched for diversifying selection

To assess whether the residues under selection correlated with specific features of kinetochore proteins, we next analyzed the structural context for our selection analyses. For this analysis, we predicted protein secondary structure using JPred4 (24) and mapped the identified residues under selection in each gene to the secondary structure predictions (e.g., coils, helices and sheets) of each kinetochore protein using the human amino acid sequence (GCF_000001405.39) as a reference (25). Enrichment analyses of structured regions across all proteins showed general enrichment of purifying selection and depletion of diversifying selection (Figure 3A; S8; S9). Most sites with evidence for episodic and pervasive diversifying selection resided in protein regions lacking secondary structure predictions (Figure S8, S9). Of all the sites in our analysis, regardless of selection type, 53.71% were located in predicted unstructured regions. Comparatively, 65.08% of episodic diversifying sites (fisher test for enrichment *p* = < 2.2e-16, odds ratio = 1.650) and 65.38% of pervasive diversifying sites (*p* = 0.0179, odds ratio = 1.630) were in an unstructured region (Figure 3A), indicating a significant enrichment for both types of diversifying selection within unstructured regions of the proteins. Accordingly, we observed a significant depletion of purifying selection sites in unstructured regions (43.82% of purifying sites, *p* = < 2.2e-16, odds ratio = 0.473) (Figure 3A; S8).

**Figure 3.**
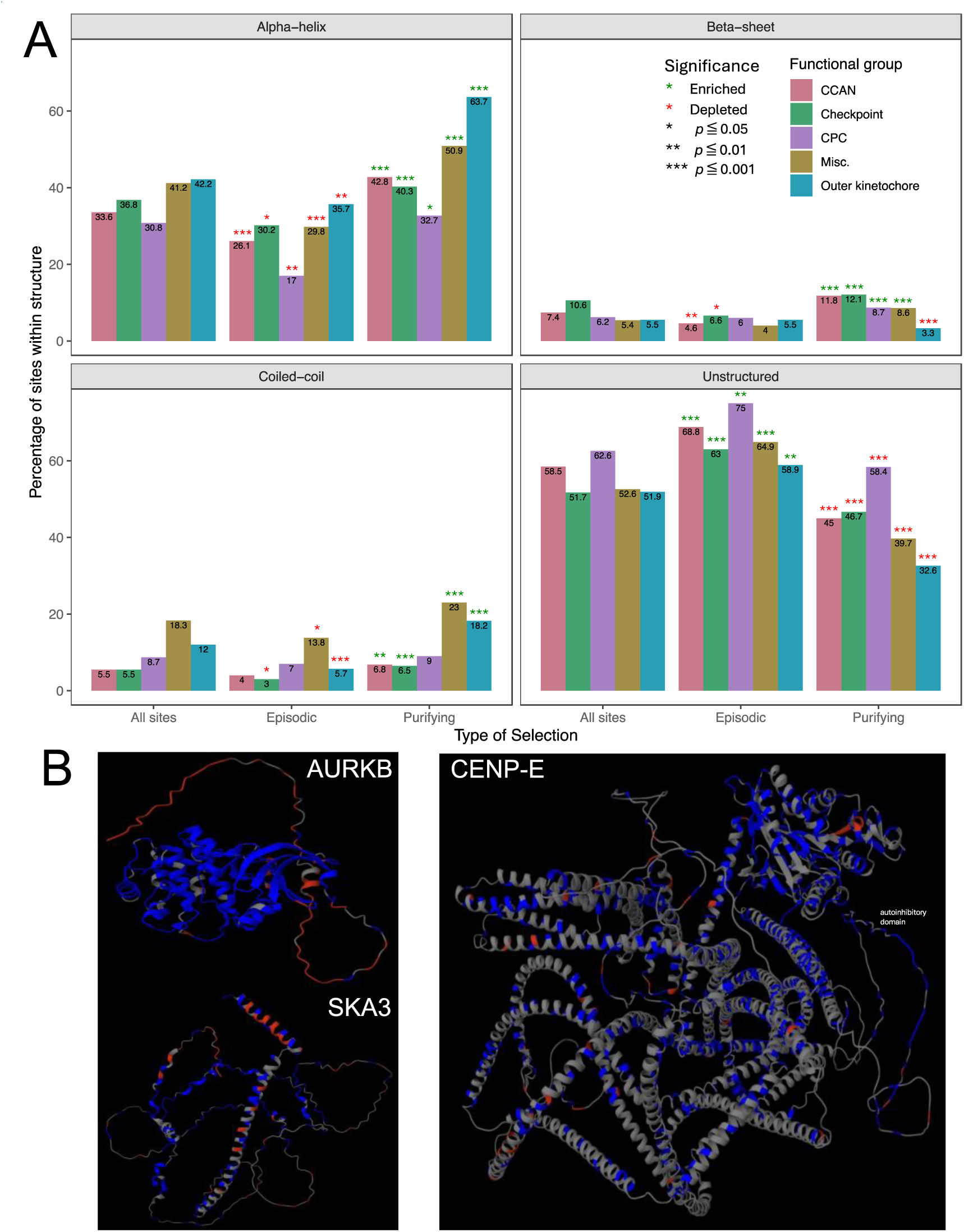
(A) Percentage of sites within each secondary structure category (alpha-helix, beta-sheet, coiled-coil and unstructured) across genes in each functional group. The bars labeled “all sites” indicate the gross percentage of sites within each type of structure regardless of whether or not the site is influenced by selection. Bars are color coded according each functional group. Significant enrichment is denoted by a green asterisk, depletion by a red asterisk. (B) AlphaFold structure predictions for AURKB (Q96GD4), SKA3 (Q8IX90), and CENP-E (Q6RT24). Sites predicted to be evolving under purifying selection are highlighted in blue; sites evolving under episodic diversifying selection are highlighted in red. AURKB exemplifies the general pattern found across all genes with enrichment of negative selection in structured regions and diversifying selection in unstructured regions. SKA3 and CENP-E exhibit contrasting patterns.

We observe the reverse when considering the various secondary structures; these regions tend to be depleted of diversifying selection and enriched for purifying selection (Figure 3A; S8). The most common secondary structure among kinetochore proteins was the alpha-helix, with 38.29% of sites located in helical regions. However, only 29.21% of episodic diversifying sites (*p* = 2.42e-16, odds ratio = 0.650) and 30.77% of pervasive diversifying sites (non sig. *p* = 0.130, odds ratio = 0.716) were in helices compared to 45.62% of purifying sites (*p* = < 2.2e-16, odds ratio = 1.785) (Figure 3A; S8).

We also annotated coiled-coil regions and beta-sheets. No significant enrichment or depletion of pervasive diversifying sites was found in either type of structure (coil *p* = 0.874, odds ratio = 0.881; sheet *p* = 0.0888, odds ratio = 0.376) (Figure S8; S9), possibly due to a lack of statistical power corresponding to the limited number of diversifying sites identified in our analysis (104 pervasive diversifying compared to 1,770 episodic diversifying sites and 14,834 purifying sites). However, we note that in both coiled-coils and beta-sheets, the percentage of pervasive diversifying sites is low (coils: 10 sites, 9.62%, sheets: 3 sites, 2.88%) with enrichment for purifying selection (coils: *p* = 4.03e-06, odds = 1.181; sheets: *p* = < 2.2e-16, odds = 2.027) and depletion for episodic diversifying selection (coils: *p* = 7.58e-09, odds = 0.594; sheets: *p* = 9.25e-05, odds = 0.659). This pattern of increased purifying selection and decreased diversifying selection in structured regions was also observed when considering the proteins in each functional group (e.g., checkpoint, CCAN, CPC, outer kinetochore and miscellaneous) with the one exception being beta-sheets in outer kinetochore proteins wherein we find a depletion of purifying selection (Figure 3A).

Contrary to this pattern, we also identified eight kinetochore proteins that are enriched for episodic diversifying selection in structured regions. Coiled-coil regions in BIRC5 (*p* = 0.036, odds ratio = 3.837), MIS18A (*p* = 0.037, odds ratio = 5.310), and OIP5 (*p* = 3.29E-04, odds ratio = 8.790) are enriched for episodically diversifying sites. Helices in OIP5 (p = 2.96E-03, odds ratio = 4.659) and SKA3 (*p* = 0.0243, odds ratio = 2.181) (Figure 3B) and sheets in ZW10 (*p* = 0.0225, odds ratio = 6.0261), SKA1 (*p* = 8.26E-04, odds ratio = 7.134), CENP-K (*p* = 0.0135, odds ratio = 9.417), and INCENP (*p* = 3.02E- 04, odds ratio = 17.494) are also enriched for episodic diversifying selection.

Furthermore, we identified selected genes that exhibited depletion of purifying selection in structured regions opposite to the general pattern of increased conservation in structured regions across all genes. The NDC80 complex subunits SPC24 (*p* = 0.0169, odds ratio = 0.407) and NUF2 (*p* = 0.0152, odds ratio = 0.565) are depleted of purifying selection in coiled-coil regions and the kinesin motor CENP- E (*p* = 3.14E-06, odds ratio = 0.634) is depleted in alpha-helices. CENP-E exhibits a contrasting pattern of enrichment of purifying selection in its unstructured regions (*p =* 1.56e-03, odds = 1.411), specifically in the unstructured tail (positions 2479 – 2575). This block of purifying selection coincides with the globular autoinhibitory domain and sequence conservation in this region could be acting to preserve its function (Figure 3B). Finally, in addition to helices being enriched for episodic diversifying selection, unstructured regions in SKA3 are depleted of episodic diversifying selection (p = 0.024, odds = 0.458) in opposition to the general trend across all genes (Figure 3B). Overall, we find that purifying selection appears to be maintaining the secondary structures of these proteins, whereas diversifying selective changes occur in unstructured regions.

### Sites in CENPC, CENPT, MIS18BP1 and TTK/Mps1 show corresponding trends in evolution with other genes among species

To explore the relationship between the evolutionary dynamics of kinetochore proteins and their molecular interactions, we calculated Spearman’s correlation coefficients (26) for each possible pair of evolving sites across all genes. We filtered the results to identify significant sites where the same three or more species evolve under episodic diversifying selection at pairs of residues, either within the same locus, or between loci. These sites, referred to as “corresponding sites”, could be indicative of compensatory evolution in which selection at a given residue precipitates corresponding diversifying selection in another residue within the same species. We used a stringent empirical Bayes factor (EBF) minimum value of 10 to denote sites under selection in a particular species, and we only considered instances of selection occurring on terminal branches representing independent selection events. We found 27 corresponding pairs of sites across all kinetochore proteins (Figure 4). 22 of those pairs were sites in two different loci, and 5 pairs were internal to the same gene. Most corresponding sites were in unstructured regions of the protein (46 of 58 sites). In the few instances where one of the sites in a pair was located in a structured region, the complementary site was in an unstructured region. The group of species under selection at both sites in a pair was highly variable and included a total of 41 different species comprising the 27 corresponding pairs (Table S2). Figure 4 depicts the network of proteins with corresponding sites.

**Figure 4.**
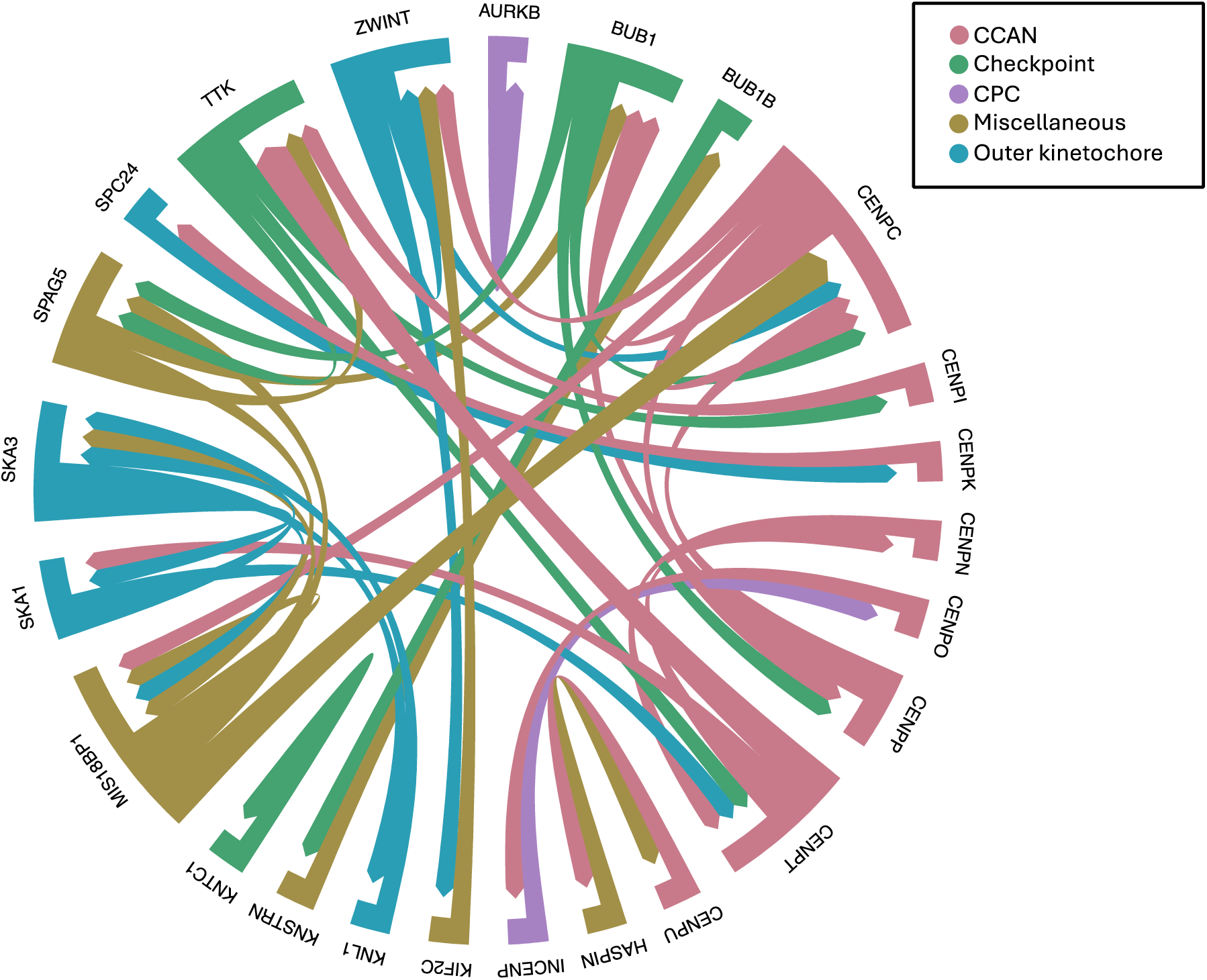
Corresponding sites represented as a chord plot. Corresponding sites were identified using a Spearman’s rho correlation and are defined as sites where the same group of three or more species are evolving under episodic diversifying selection. The color scheme corresponds to the functional grouping as described in Figure 1. Arrows point to the gene that a site corresponds with. The width of the arrow is representative of the number of sites in the gene from which the arrow came corresponding to a site in the interacting partner. For example, the thick red arrow from CENPT pointing to TTK/Mps1 represents two sites within CENPT corresponding to the same site in TTK/Mps1.

The four genes with the highest number of corresponding sites with other genes in the analysis are CENPC, CENPT, MIS18BP1, and TTK/Mps1 (Figure 4). Four sites within CENPC have corresponding evolutionary patterns with five sites in four other genes (one site in BUB1, one in CENPP, two in MIS18BP1, and one in ZWINT) and one pair of internally corresponding sites. CENPT has four sites with corresponding evolutionary patterns to three sites in three other genes (one site in each of CENPN, SKA1, and TTK/Mps1), MIS18BP1 has four sites with corresponding patterns to three sites in three other genes (one site in each of CENPC, SKA3, and SPAG5/Astrin) and one pair of corresponding internal sites. Finally, TTK/Mps1 has three sites with corresponding patterns to four sites in three other genes (one site in CENPI, one site in SPAG5/Astrin, and two sites in CENPT) (Figure 4). There were six other genes in the analysis with more than one non-internal correspondence with other genes (BUB1: three, SKA3: three, SPAG5/Astrin: three, CENPP: two, SKA1: two, and ZWINT: two) (Figure 4).

Our ability to detect sites undergoing corresponding evolutionary changes could be biased by the degree of selection among loci. However, we observe little overlap between the genes with the highest number of sites under episodic diversifying selection and the genes with the most corresponding sites. For example, CENPE had the third highest number of sites under episodic diversifying selection, but has no evolutionary correspondences, internal or otherwise. Even when taking into account both the number of episodic diversifying sites and the number of species’ sequences included in each gene’s alignment (applying the logic that two genes must have species in common for there to be correspondences), these factors explain only a small amount of variation in the number of correspondences (*p* = 0.004, adj. R^2^ = 0.125). We are therefore encouraged that our analyses reflect real evolutionary dynamics between genes. In summary, we were able to identify 27 pairs of sites where the same group of species independently experience diversifying selection, potentially indicative of compensatory evolution. The genes that stand out as having the highest number of these sites and therefore could be considered “hotspots” of correlated selective change are CENPC, CENPT, MIS18BP1 and TTK/Mps1.

### Genes with sites exhibiting corresponding trends in evolution experience little purifying selection

In an effort to further understand the evolutionary correspondences described above, we compared the number of corresponding sites with the percentage of sites under purifying selection within each gene. We observed that the genes with the most corresponding sites experience little purifying selection when compared to other genes in their functional grouping (Figure S10). For example, CENPC has the highest number of sites with corresponding patterns (five corresponding sites with other loci) (Figure 4). The percentage of sites under episodic diversifying selection (10.29%) for this gene falls within the average for CCAN genes (CI: 8.69 - 12.87%). However, CENPC has a below average percent of sites under purifying selection (14.21%) when compared to the other CCAN genes (CI: 34.80 - 55.00%). Out of the ten genes with two or more sites corresponding with other genes, eight experience a level of purifying selection that is significantly below the average for the other genes in their functional group (Figure S10). This suggests that a relaxation of purifying selection may be associated with the evolution of correspondent sites.

## DISCUSSION

Here, we used the rapid evolution of the kinetochore as a model to analyze the relationship between evolutionary patterns and macromolecular protein complex assembly and function. Our approach interrogated three levels of evolutionary dynamics including clade-specific trends, gene-specific trends, and coordinated evolution between genes.

### Lineage specific trends

In contrast to previous work, which found that evolutionary pressure among kinetochore loci was consistent across mammals (20), our alternative methodology utilizing branch and site information found that the CCAN genes of Afrotherians are evolving significantly more rapidly than other orders (Figure 1B). Our approach provided a robust and sensitized method for the evaluation of evolutionary pressures, potentially defining changes that had not been detected in previous work. However, Afrotheria experience a level of selection within checkpoint genes that is consistent with other mammals, highlighting the tight conservation of these proteins even in species where other kinetochore genes evolve rapidly. Afrotheria is also the most morphologically diverse clade of mammals (27), including species such as the lesser Madagascar tenrec (*Echinops telfairi*), which grows to between 5.5 and 7 inches (28), and the African elephant (*Loxodonta africana*), the largest extant terrestrial animal (29). Our finding that this order is experiencing a high amount of diversifying selection is interesting in that the diversity of body plans appears to be reflected in the sub-cellular composition as well.

### Functional module trends

In line with recent findings (20), we found that the genes that form the constitutive centromere associated network (CCAN) and the outer kinetochore genes experience significant diversifying selection, especially in comparison to the checkpoint genes (Figure 2; S3). The CCAN genes particularly stand out in analyses of pervasive diversifying selection (Figure S2; S3; S4B; S6). When considering the potential drivers of this rapid evolution, it is worth noting that centromeric loci comprise some of the most rapidly evolving DNA sequences in the genome (30). Many of the CCAN proteins either interact with DNA directly or are closely proximal to the DNA (13, 31–34). Although the centromere is primarily specified epigenetically based on the presence of the centromere-specific histone H3 variant CENPA (11), recent work has highlighted contributions of DNA sequence features in centromere function (30, 35, 36). Thus, these changes in CCAN proteins across species may allow cells to adjust and compensate for changes to the underlying DNA.

Another evolutionary force potentially shaping kinetochore function is brain development. The cell division machinery is expressed universally. However, the most common phenotypes in humans resulting from kinetochore gene mutations involve the brain, which requires many well controlled cell divisions for proper development. Mutations in kinetochore genes including KNL1 (37), CENPT (38), CENPE (39), and BUB1 (40) among others have been linked to microcephaly where a reduction in the number of neurons in the developing brain leads to a smaller than normal head size (41). A recent study looking at a ubiquitously expressed mutation in KNL1 found that it only produced abnormalities in neural progenitor cells and not in fibroblasts or neural crest cells (37) highlighting the unique role of kinetochore function in the development of the brain. Therefore, differences in brain development across mammals could create evolutionary pressures that would help to explain our findings of heightened diversifying selection in this structure.

When considering the evolutionary pressures acting on the kinetochore, we must also acknowledge that it must function properly during two inherently different processes, mitosis and meiosis. In contrast to mitosis, meiotic cells must undergo two consecutive rounds of DNA segregation to generate gametes containing half the number of chromosomes of the original parent cell. To accommodate this, cells must alter how the DNA is organized in the cell and consequently the interactions between the kinetochore and spindle microtubules must also change. The Shugoshin protein family exemplifies the different requirements of these two processes. In mitotic cells, the cohesin complex mediates attachments of sister chromatids until anaphase onset, where it is degraded allowing chromatids to separate (42, 43). During this process, Shugoshin 1 (SGO1) acts to protect kinetochore localized cohesin from premature removal (44). In meiosis, this process must be altered to allow segregation of homologous chromosomes in meiosis I, while maintaining cohesion between sister kinetochores. To achieve this, germ cells rely on a second Shugoshin ortholog, Shugoshin 2 (SGO2) that protects sister chromatid cohesion during anaphase I (43, 45). In humans, loss of SGO2 contributes to age-related aneuploidy in oocytes (46). In contrast, loss of SGO2 during mitosis does not affect sister chromatid cohesion. Instead, during mitosis SGO2 plays an entirely different role ensuring proper kinetochore-microtubule attachment (47, 48). SGO2 was previously identified as experiencing a high amount of diversifying selection compared to other kinetochore genes (20). Although we did identify a depletion of purifying selection in SGO2, it did not stand out as experiencing substantial diversifying selection in our analyses. However, it is worth noting that SGO2 had the largest percentage of sequences removed due to the presence of unalignable regions, which could suggest a highly variable exon structure between species or regions of rapid evolution. This is surprising given its critical role in meiosis, but a lack of negative selection may be indicative of the flexibility required to function in two different roles.

Another protein that stands out as having a lower number of sites under purifying selection than expected is the checkpoint protein BUB1, which is partially responsible for the recruitment of SGO2 to the inner centromere (46, 47, 49). In contrast to CCAN and outer kinetochore genes, checkpoint genes experience low diversifying selection and high purifying selection. In line with this finding, mutations in checkpoint genes are often poorly tolerated by cells and can predispose cells to cancer (50, 51). However, when compared to other checkpoint genes, BUB1 has the lowest percentage of sites under purifying selection and higher diversifying selection (Figure 2; S4). As mentioned above, Biallelic BUB1 mutations have been shown to cause microcephaly and other developmental delays (40), but patient cells with these mutations still exhibited a functioning Spindle Assembly Checkpoint (SAC). Unlike most of the other checkpoint proteins, some functions carried out by BUB1 can also be performed by other proteins, likely affording BUB1 a tolerance to sequence changes that is not afforded to other proteins which do not have a “backup” mechanism. For example, BUB1 and the CENPO complex act in parallel to recruit the mitotic kinase Plk1 to kinetochores such that these complexes can partially compensate for each other (52).

Finally, the elevated rate of evolution observed among kinetochore loci may also be influenced by meiotic drive, the phenomena in which selfish genetic elements act to increase segregation of a particular chromosome homolog to the haploid egg (53). Unlike mitosis, in which all chromosome pairs are transmitted to offspring, female meiosis results in only a single viable egg product in which one quarter of chromosomal complement is successfully passed while the remaining genetic material is evicted from the oocyte into the polar bodies. In this asymmetric cellular division, proteins that promote stronger kinetochore microtubule attachments, or those that could influence orientation of the chromosomes, may be more likely to be transmitted to the single egg, allowing for non-mendelian segregation and a strong source for evolutionary insight (53–56).

### Selection in the context of protein secondary structure

In our analyses, diversifying sites were significantly enriched in regions lacking clear secondary structure predictions and depleted in regions with predicted secondary structure (Figure 3A). The opposite was true for purifying selection, which appears to be acting to maintain the secondary structure of the protein while minimizing constraints in unstructured regions. Previous genome-wide studies found similar results, where sites that form disordered regions of proteins are more likely to be under diversifying selection (6) than structured sites. A more recent study (57) including the genomes of 90 mammalian species found that increased diversifying selection, coupled with a relaxation of purifying selection, contributes to heightened evolutionary rates in unstructured regions. Additionally, this study found that protein interaction networks comprised of disordered proteins formed larger interacting clusters (over 50 proteins) than clusters formed by ordered proteins (< 8 interacting partners). We hypothesize that the structural and evolutionary flexibility afforded to proteins by disordered regions facilitates the maintenance of a multitude of functional relationships (e.g., protein-protein interactions) with other proteins. Many kinetochore interaction surfaces exist within regions lacking clear secondary structure, such as for the short sequences in CENP-C that interact with the MIS12 complex (58, 59) or in CENP-T for its interactions with the NDC80 complex (14, 60, 61). Thus, evolutionary changes in such regions may preserve overall secondary structure and protein folding, while allowing changes in short linear interaction motifs (SLiMs), presenting an avenue by which evolution can operate within a cellular protein complex and larger protein networks.

### Corresponding sites reside in functionally related genes

We propose that the rapid evolution exhibited by kinetochore loci is due in part to compensatory changes between loci. Changes in a given kinetochore protein may necessitate changes in another protein to maintain the function of the higher order structure. Support for this hypothesis is given by previous analyses showing that functionally-interdependent kinetochore proteins tend be lost or gained together across eukaryotes, suggesting co-evolution between kinetochore subcomplexes (62). We hypothesize that, if compensatory changes are responsible for some of the variation in kinetochore genes, then some sites between proteins should exhibit similar patterns of evolution. We searched for these evolutionary correspondences in our data by identifying sites with correlated evolution using a Spearman rank correlation test (Figure 4). Among the genes with the highest number of correspondences with sites in other genes are the “linker” proteins CENP-C and CENP-T. CENP-C and CENP-T represent two avenues for connecting the inner kinetochore to the outer kinetochore and these are the core regions where the inner kinetochore associates with centromeric DNA (63), with centromere DNA sequences also displaying rapid evolution (30). Acting as bridges, CENP-C and CENP-T must adapt to maintain their relationship with centromeric DNA, which could necessitate downstream changes to sustain their connection to the outer kinetochore.

MIS18BP1 and TTK/Mps1 also show multiple correspondences (Figure 4). Two sites in MIS18BP1 correlate with a site in CENP-C, which recruits MIS18BP1 to the centromere for CENP-A deposition (64). Two sites in CENP-T correspond with a site in TTK/Mps1, which modulates the recruitment of CENP-T in *Saccharomyces cerevisiae* (65). Thus, our findings could reflect these established functional relationships between the corresponding proteins. On the structural level, most of the corresponding sites are found in unstructured regions, in line with our finding that most sites under diversifying selection are located in these parts of the proteins. Taken together, our data suggest that compensatory evolution that maintains functional connections between proteins, localized largely to unstructured regions, is an important feature of the evolutionary dynamics of the kinetochore. In this way, the evolution of the kinetochore is dictated by something akin to a macromolecular Red Queen process (66), where individual proteins must perpetually adapt to changes in their direct or indirect functional partners to maintain a critical cellular function.

## CONCLUSIONS

Our analysis of the kinetochore complex reveals lineage, protein interaction module, and gene-specific evolutionary dynamics that highlight putative routes by which selection can act upon a critical macromolecular complex without perturbing its function. First, we showed that Afrotheria, the most morphologically disparate mammalian order, exhibits elevated rates of kinetochore protein evolution compared to other mammalian orders, suggesting the possibility that diversifying selection in the kinetochore is involved with morphological diversity. Next, we also observed a selection differential on the level of individual proteins and their interaction modules. The CCAN and outer kinetochore proteins display significantly greater diversifying selection than those of other modules, whereas the checkpoint proteins are held tightly in check by purifying selection. We further show that much of the selection detected across kinetochore proteins is localized to unstructured regions, providing a partial explanation for how protein interactions essential to the cell may be maintained while the larger complex is molded by selection. Finally, consistent with this hypothesis, we show that a subset of residues under diversifying selection show significant evolutionary correspondence between species. Our evolutionary insights could inform future functional analyses of the kinetochore and more broadly guide our understanding of how macromolecular complexes that are essential to cellular function may be molded by selection pressure.

## METHODS

### Genome inclusion and sequence processing

Genomic protein sequences from 70 mammalian species and four outgroups were downloaded from NCBI’s Reference Sequence Database (RefSeq) (Table S1) (25). We considered genome quality as well as taxonomic breadth and completeness when deciding which species to include. The final dataset included representatives from 15 mammalian orders: 13 Artiodactyla, 12 Carnivora, 12 Primates, 11 Rodentia, 6 Afrotheria, 3 Chiroptera, 2 Diprotodontia, 2 Lagomorpha, 2 Perissodactyla, 2 Cingulata, and one each of Eulipotyphla, Scandentia, Pholidota, Dasyuromorphia, and Didelphimorphia (Figure 1B). The protein sequences were given to OrthoFinder (v2.3.3; t=48) (67) and organized into 27,192 orthogroups. The orthogroups containing the human sequences for each of the kinetochore genes of interest were identified. We chose to analyze 57 kinetochore genes which represent a comprehensive view of the structure and function of the whole complex, while also eliminating concerns of pleiotropy which would complicate the interpretation of results by including genes that solely contribute to the process of cell division (Figure 1A). In three cases, human sequences for the same gene were sorted into more than one orthogroup (AURKB, CENPM and SKA2). In each case, the orthogroup with the largest number of sequences was chosen to represent that gene. Nucleotide sequences were required to run downstream evolutionary analyses, so a custom script was written to query the NCBI database (68) and retrieve the corresponding nucleotide sequences encoding the proteins. In a few instances the nucleotide sequence for the wild boar (*Sus scrofa*) had been removed from the database due to standard annotation processing and were withdrawn from our analysis. CDS data from the nucleotide entry on NCBI was fed into another custom script which trimmed each sequence to only include the protein-coding region. The script also removed the stop codons at the end of the sequence and removed any sequences with in-frame stop codons.

For genes in which multiple splice variants were present within a species, CD-HIT-EST (v4.7; c=0.98, n=5, d=0) (69) was run on the sequences at a level of 98% similarity clustering sequences with a 2% or less difference. Following this step, we ensured that each orthogroup contained only one human sequence, representing the most highly expressed transcript in human cells that generates the full length annotated protein or, when not possible, maximizes sequence length (Table S3).

### Sequence alignment and gene trees

The sequences for each gene were aligned twice, first with MAFFT v.7.305b (70) using the L-INS-I option. The alignments were visualized using SeaView v.5.0.4 (71) and edited to remove sequences with unalignable regions (gaps larger than ∼20 bps). The pruned alignments were re-aligned using a codon-aware aligner, MACSE (v2.05; prog=alignSequences) (72). The frameshift symbol (“!”) was removed to accommodate the requirements of downstream evolutionary analyses, then the alignments were given to iqtree (v1.6.12, m=GTR) (73) for gene tree generation using a generalized time reversible model. The trees were rooted with the vertebrate outgroup species. For each gene, the nucleotide alignment and the gene tree were concatenated into one file that would be used as input for sequence analysis. In order to help with interpretation of results, only sites where the human sequence had a residue present were considered in all further analyses.

### Selection analyses for inferring sites and branches under selection

Software from the hyphy (v2.5.26) (74) suite of analyses was used to infer sites evolving under diversifying and purifying selection and branches under diversifying selection for all 57 genes. aBSREL v2.2 (21) was used to detect branches evolving under episodic diversifying selection across all 57 genes. As was the case for all hyphy analyses used, the software was given the nucleotide alignment and gene tree as input. Significance for aBSREL was evaluated using a Likelihood Ratio Test at a threshold of p <= 0.05.

Sites under episodic diversifying selection were identified according to MEME v2.1.2 (22) using a p-value cutoff of 0.1 and testing for selection across all branches. Empirical Bayes factor (EBF) values generated by MEME were used to identify the branches where selection at each episodic diversifying site is most likely to be occurring. At each episodic diversifying site, we hypothesized that a certain species was evolving if the EBF value was 10 or larger on the terminal species branch. We made the decision to ignore evidence of selection occurring on internal branches, because it would be difficult to draw conclusions about which species that selection event is influencing. Sites displaying evidence of pervasive diversifying selection and purifying selection were identified using FUBAR v2.2 (23) using a posterior probability threshold of 0.9.

### Statistical analysis

To test for differences in the mean number of sites under selection per gene across mammalian clades, an ANOVA was conducted, followed by a Tukey comparison (alpha = 0.05) of least squares means (CI = 0.95) for a significant ANOVA using the lsmeans v.3.30-0 package (75) in R v.4.2.2 (76). We only considered orders with at least three representative species included in our analysis, and we tested each functional group of genes separately in order to detect functional group trends. Differences due to between gene variance were accounted for in the model, as well as any significant interaction between the gene and the order. This same workflow was employed to test for differences in the mean number of genes under selection summed for each functional category across orders according to aBSREL, as well as for testing for differences in the mean percentage of sites under selection according to functional group.

Relationships between variables in our study, such as the relationship between the number of sites under selection and the length of a gene, were detected using a Pearson’s correlation at a p-value cut-off of 0.05. All tests for enrichment were conducted using a Fisher’s exact test. All analyses were implemented in R v.4.2.2.

### Secondary structure prediction and enrichment analyses

Secondary structure was predicted using Jpred4 (24) given the amino acid sequence of the human protein for each gene. Results from the selection analyses were combined with the results from JPred4, and enrichment or depletion of sites with evidence of selection in coils, helices and sheets was detected using a Fisher’s exact test implemented in R v.4.2.2. The same test was used to test for enrichment or depletion in unstructured regions defined as sites not residing in a coil, helix or sheet according to the structure results.

### Detecting corresponding evolutionary patterns

To highlight sites that could potentially be evolving in a correlated manner, we calculated a Spearman’s rho coefficient using the Hmisc (v.4.7-2) (26) package in R (v.4.2.2) for all pairs of evolving sites using a rho cutoff of 0.9 and a false discovery rate (FDR) cutoff of 0.01. For each gene, the analysis was given a matrix of 1 or 0 values for each site/species combination, where a 1 represented a site with an EBF value of 10 or above on that particular species branch and a 0 represented a value below 10. EBF values were extracted from the output of MEME, which assigns an EBF value for each branch/site combination representing the likelihood that a particular branch is undergoing selection at a specific site. Significant pairs were filtered to identify pairs of episodically diversifying sites where the same three or more species have an EBF value of 10 or above.

## ACKNOWLEDGEMENTS AND FUNDING SOURCES

We thank Lindy McKee and Stephen Mrenna for their research assistance, as well as Toni Westbrook for providing technical support. Thank you to the Plachetzki and MacManes lab members for their helpful feedback on the manuscript. This work was supported by the NSF (grant 2029868 to I.M.C, grant 1755337 to D.C.P.) and a Damon Runyon postdoctoral fellowship to A.L.N.

