## Supplemental Figures and Tables for "Structural constraints and drivers of molecular evolution in a macromolecular complex; the kinetochore"

### SUPPLEMENTAL DATA

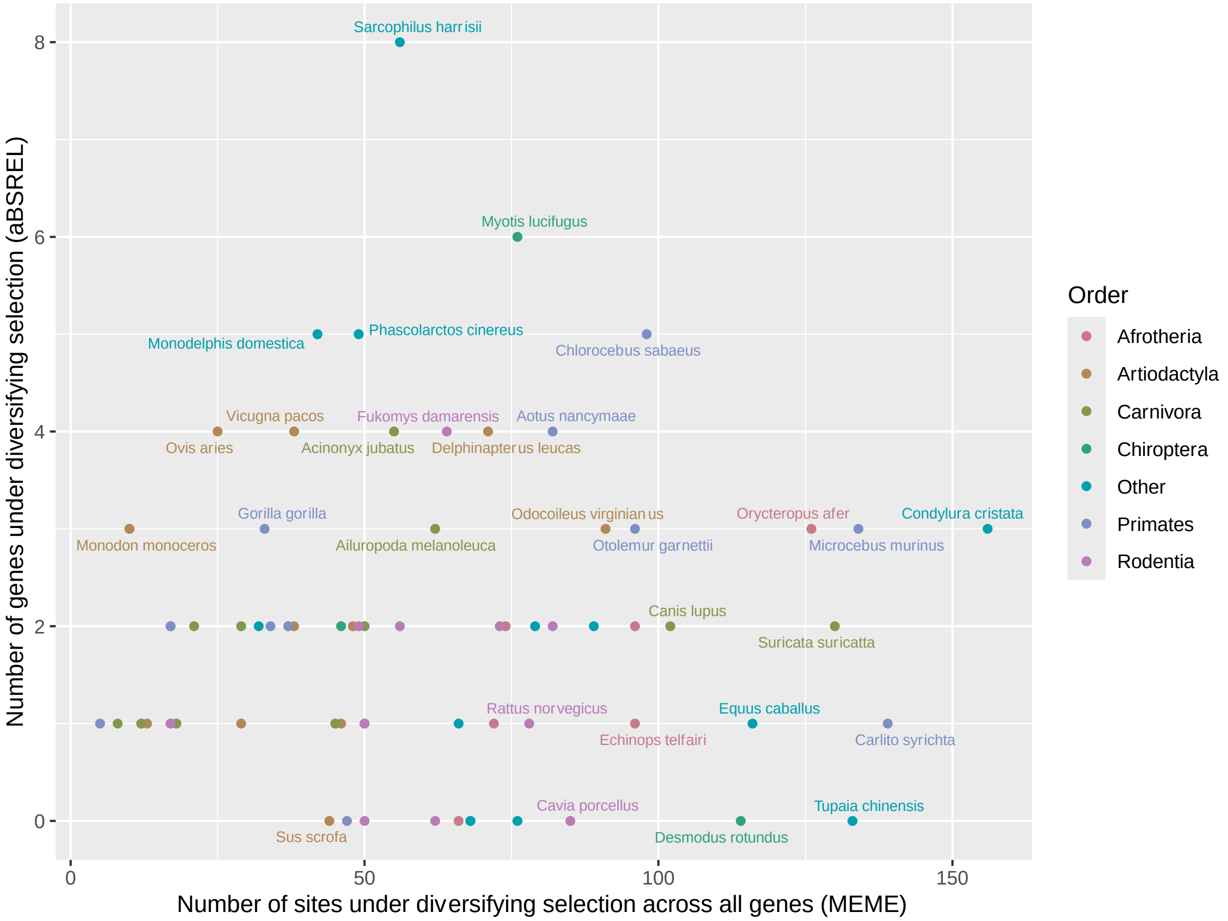


**Figure S1.** Comparing the degree of diversifying selection in each species according to aBSREL and MEME. MEME identifies the likelihood that a site is evolving under episodic diversifying selection on a specific branch. The number of sites across all kinetochore genes likely to be under selection on the branch leading to the species lineage is plotted on the x-axis. aBSREL does not test individual sites; instead the number of genes found to be under selection for a given terminal species branch is plotted on the y-axis. Species are color-coded according to the order for which they belong. Species for which their order had less than three representatives are grouped into a category called “Other”.

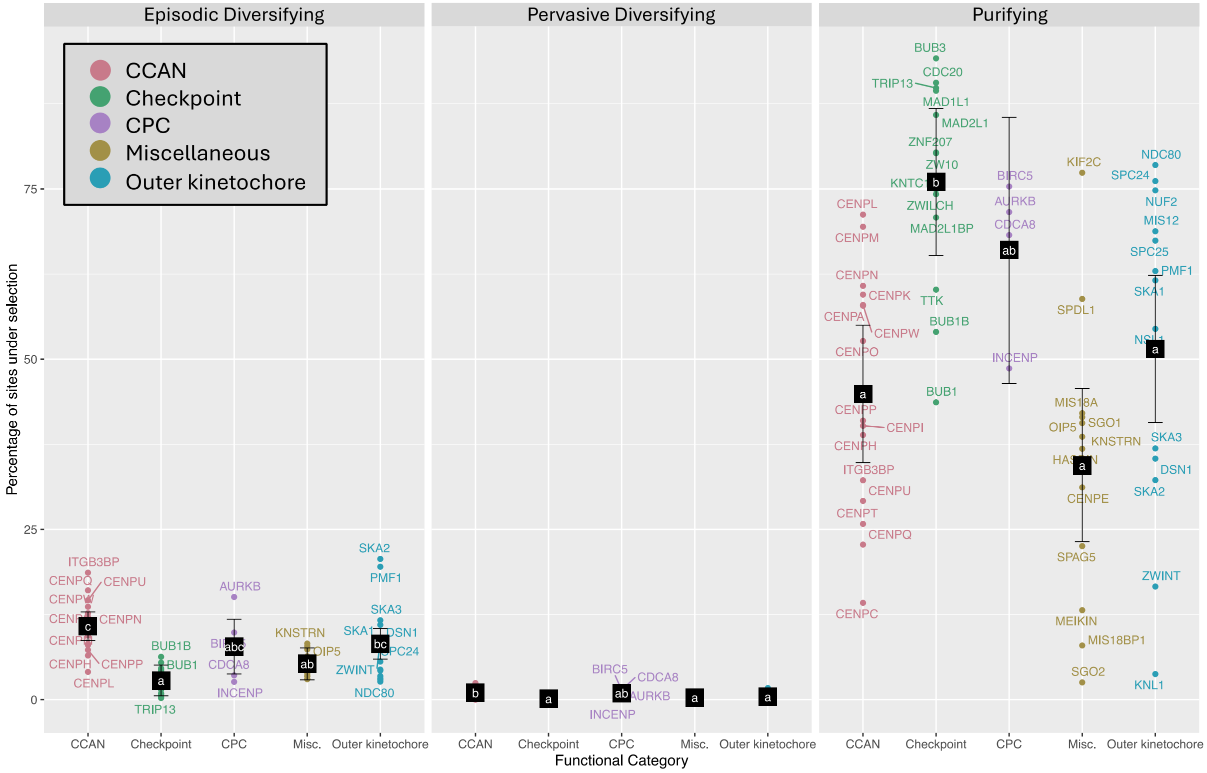


**Figure S2.** Extension of figure 2 to include pervasive diversifying selection. The percentage of sites under selection in each gene according to functional group (red: CCAN genes, green: checkpoint, purple: CPC, gold: miscellaneous and blue: outer kinetochore). Black squares are the least squares means for each functional category and bars are the 95% confidence interval around the means. Letters represent significance groups according to a pairwise Tukey test comparing the means of all five groups.

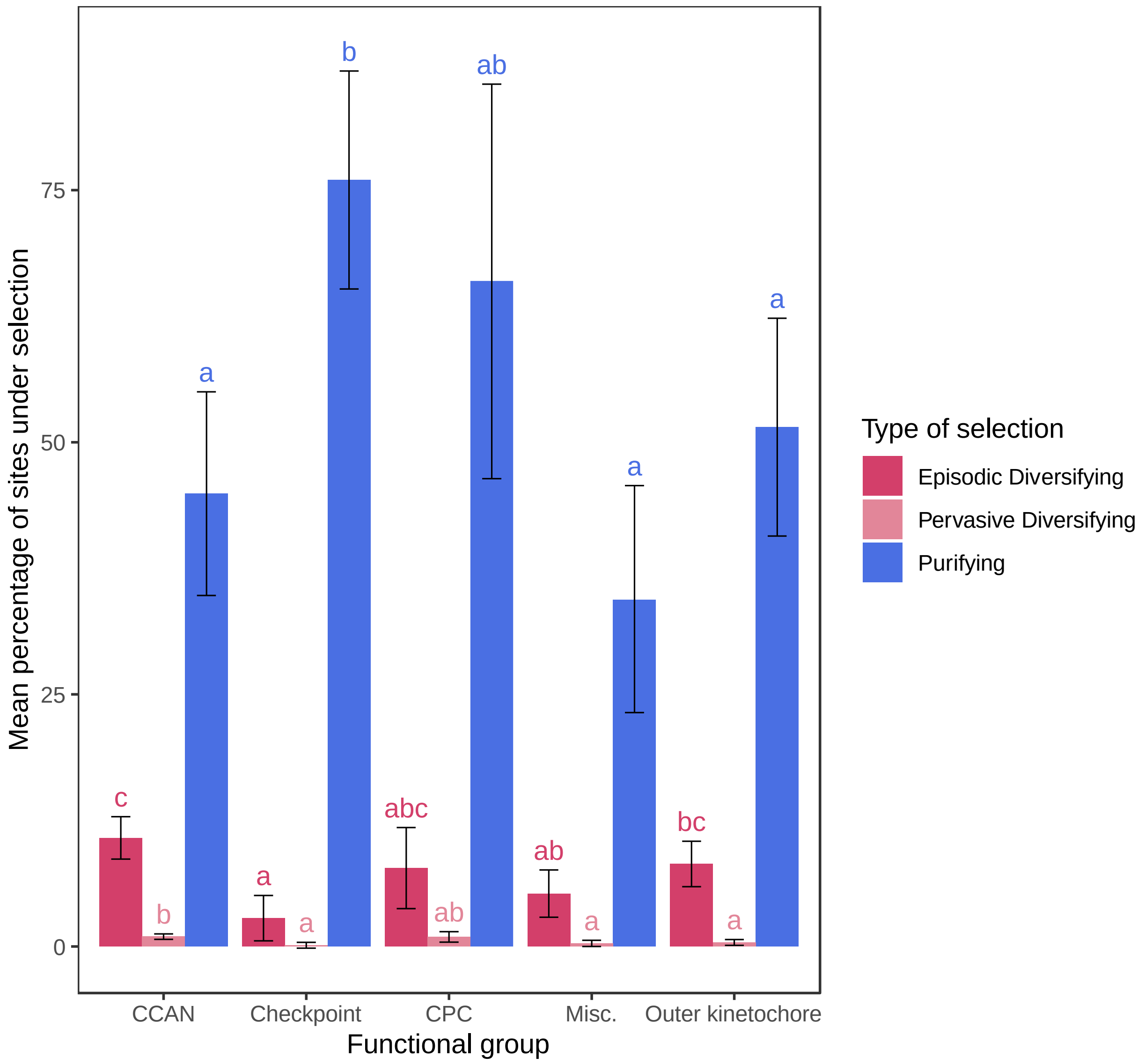


**Figure S3.** Mean percentage of sites under selection (y-axis) in each functional group (x-axis). Colored bars denote the three types of selection (dark red: episodic diversifying, light red: pervasive diversifying, blue: purifying). Letters are significance groups according to a Tukey test comparing the least squares means of the five functional groups. Error bars are 95% confidence intervals around the mean.

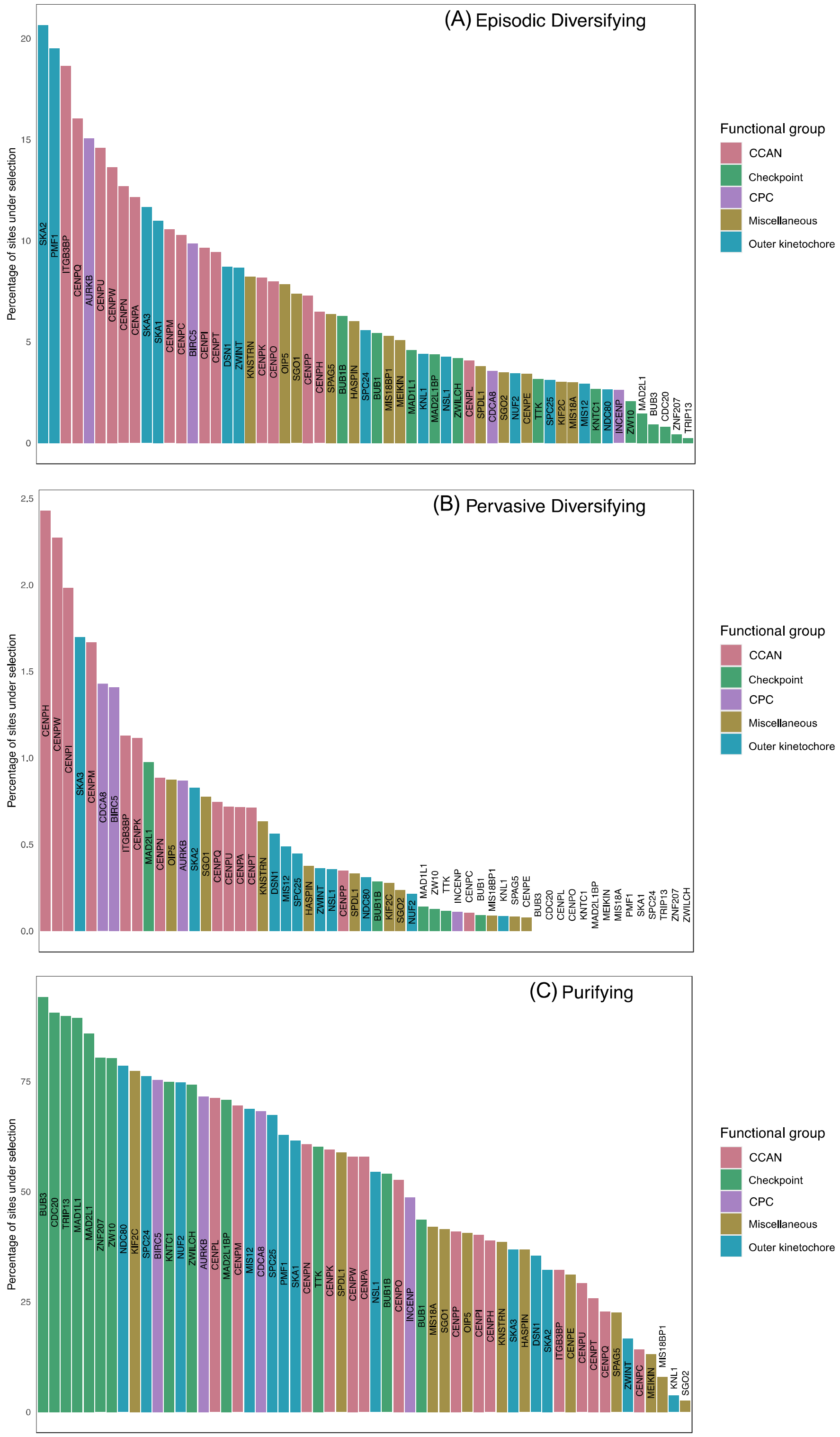


**Figure S4.** Genes ordered from highest to lowest percentage of sites under (A) episodic diversifying (B) pervasive diversifying and (C) purifying selection and colored according to functional grouping.

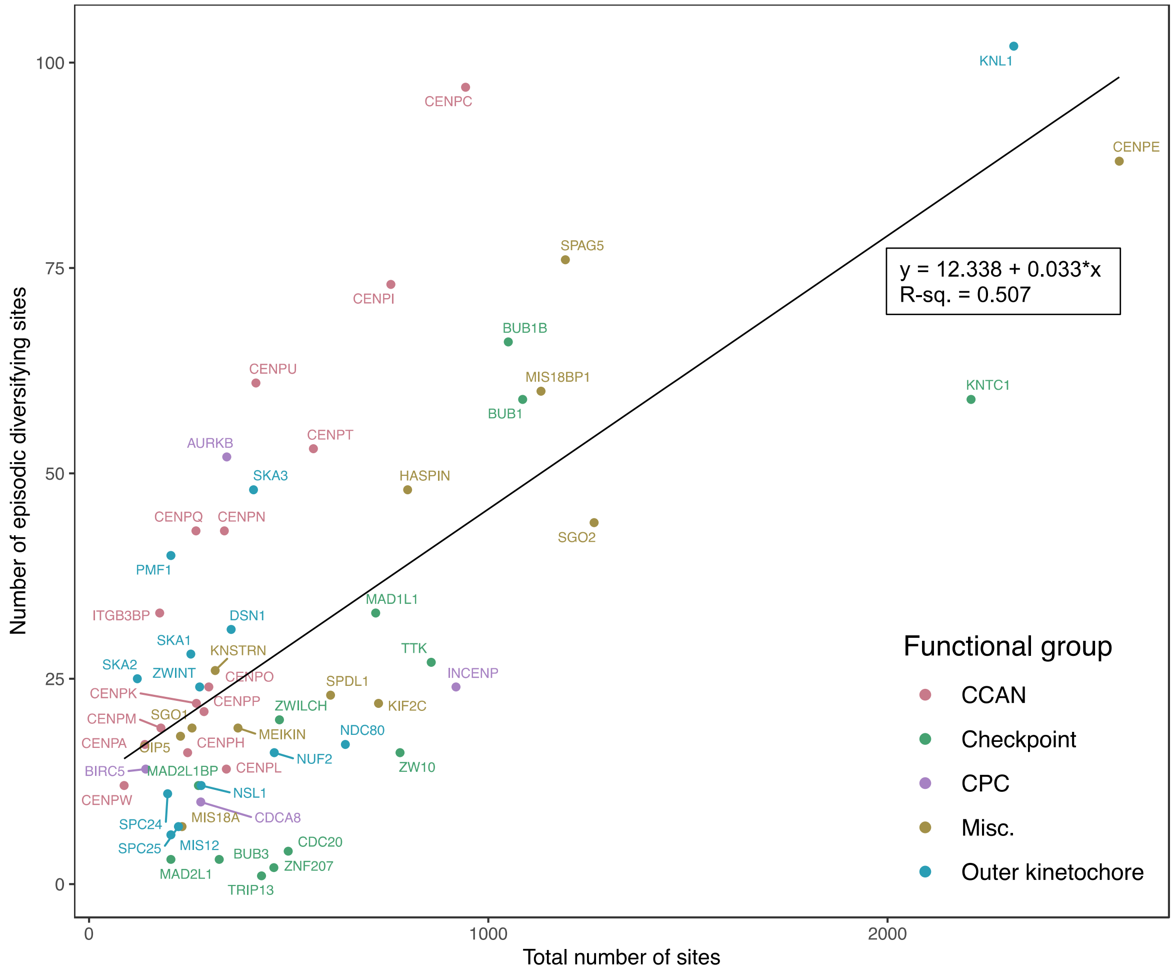


**Figure S5.** The length of each gene (x-axis) plotted against the number of sites with evidence of episodic diversifying selection according to MEME (y-axis). The trendline shows a significant positive linear relationship between the two variables ($p$ = 5.164e-10, $R^2$ = 0.5074). Genes are colored according to functional group.

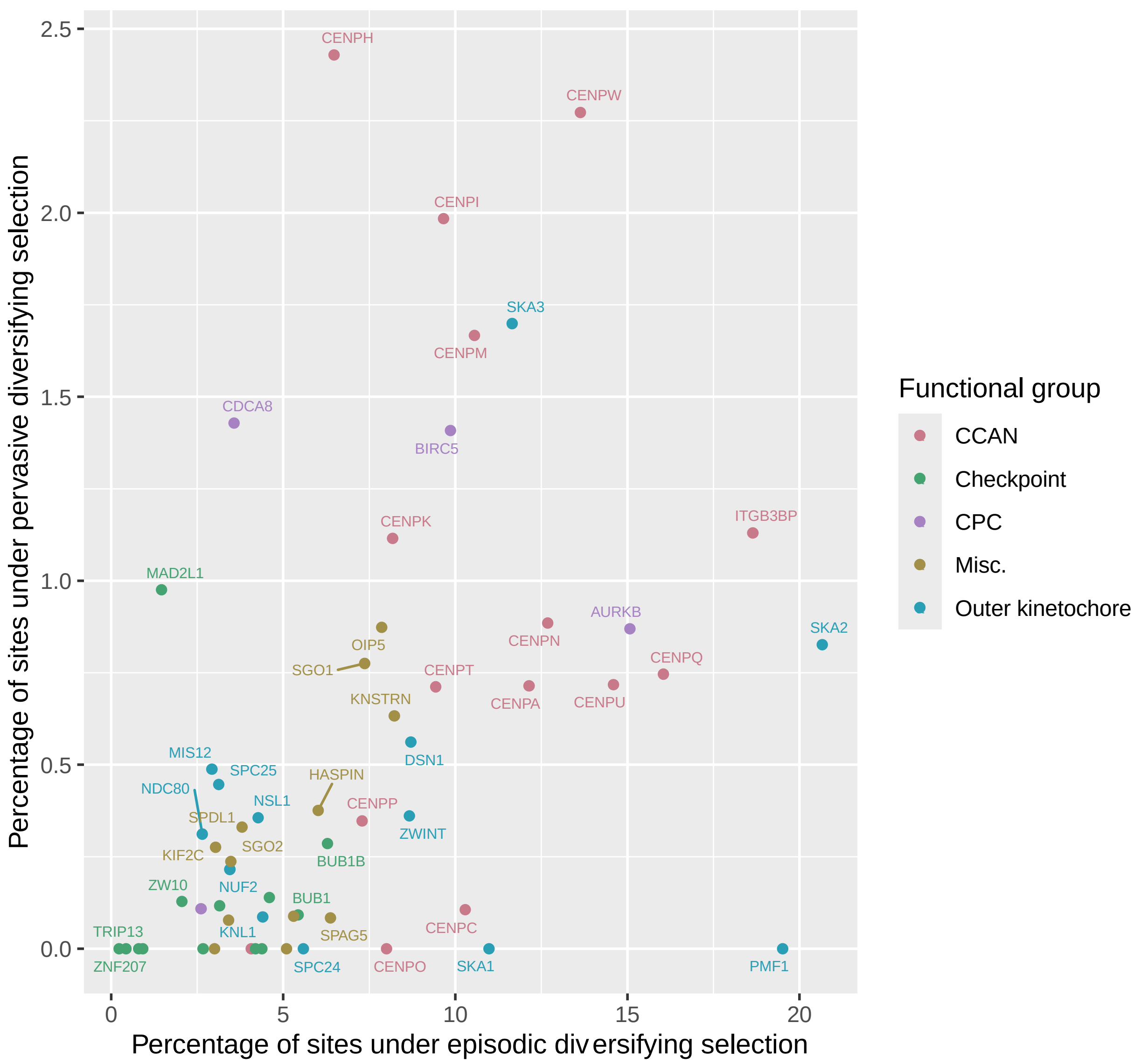


**Figure S6.** Degree of episodic versus pervasive diversifying selection for each gene colored by functional grouping. The x-axis is the percentage of sites evolving under episodic diversifying selection (ranging from 0 - 21%); the y-axis is the percentage of sites evolving under pervasive diversifying selection (ranging from 0 - 2.5%).

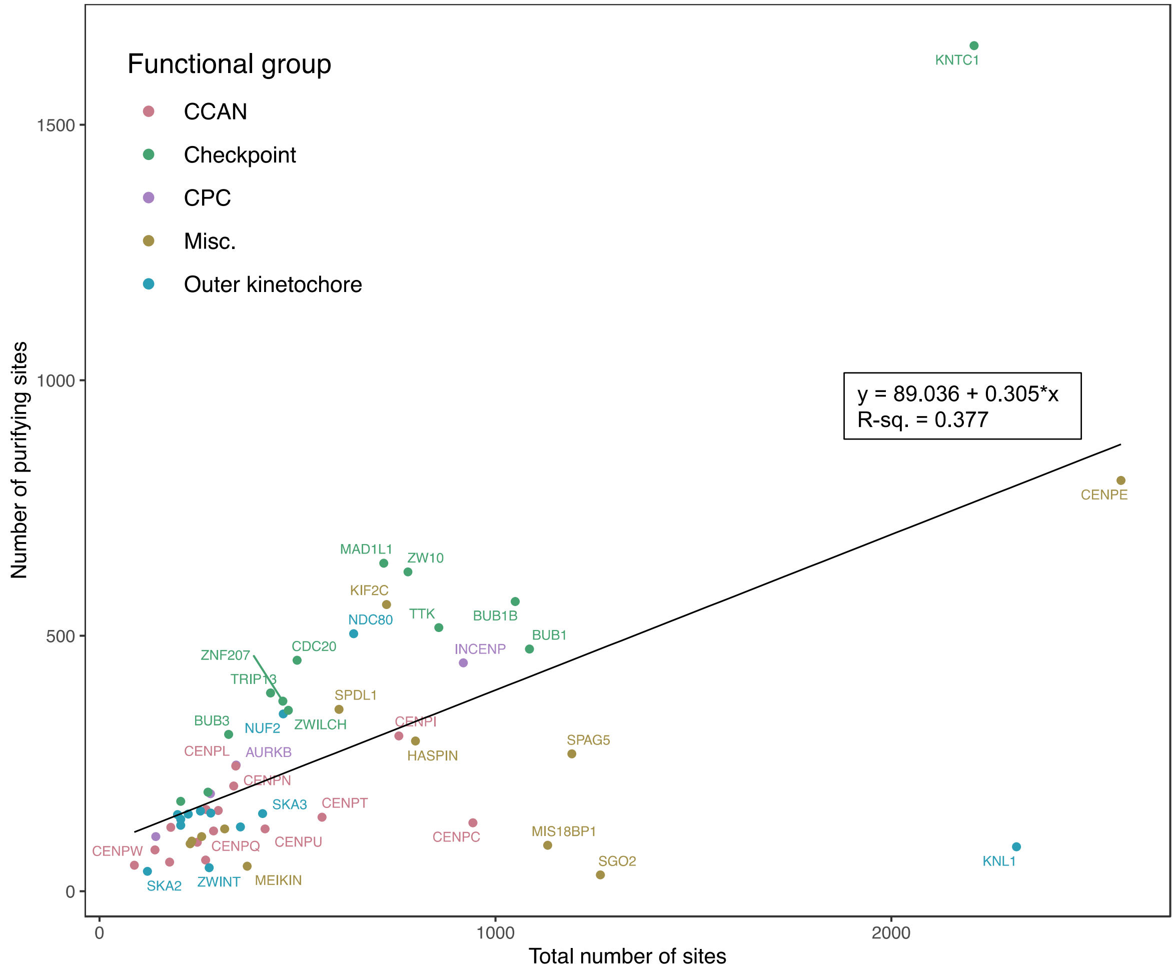


**Figure S7.** The length of each gene (x-axis) plotted against the number of sites with evidence of purifying selection (y-axis). The trendline shows a significant positive linear relationship between the two variables ($p$ = 3.802e-07, $R^2$ = 0.377). Genes are colored according to functional group.

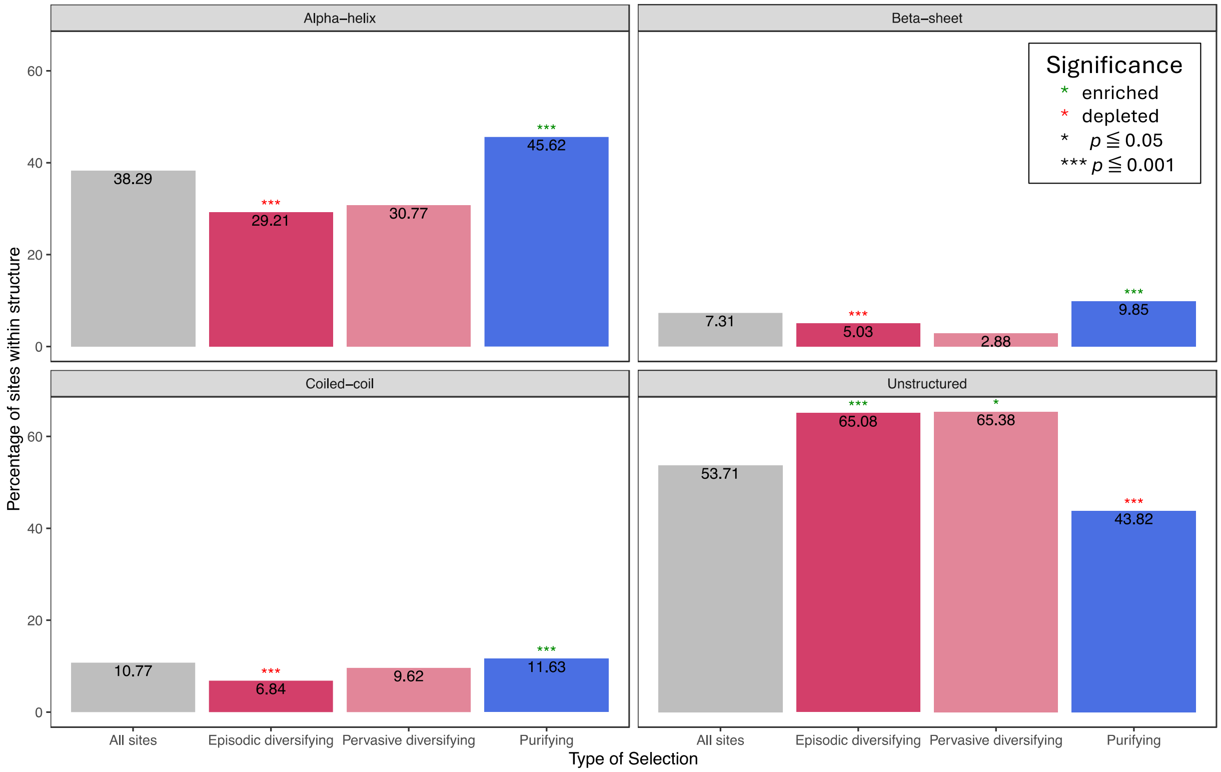


**Figure S8.** Percentage of sites within each secondary structure region (alpha-helix, beta-sheet, coiled-coil and unstructured) combined across all genes in the analysis. On the x-axis is the type of selection (episodic diversifying, pervasive diversifying, or purifying) and the "all sites" group is the percentage of sites within the particular structure region regardless of selection. Significant enrichment of selection type within a particular structure is denoted with a green asterisk; depletion is denoted with a red asterisk.

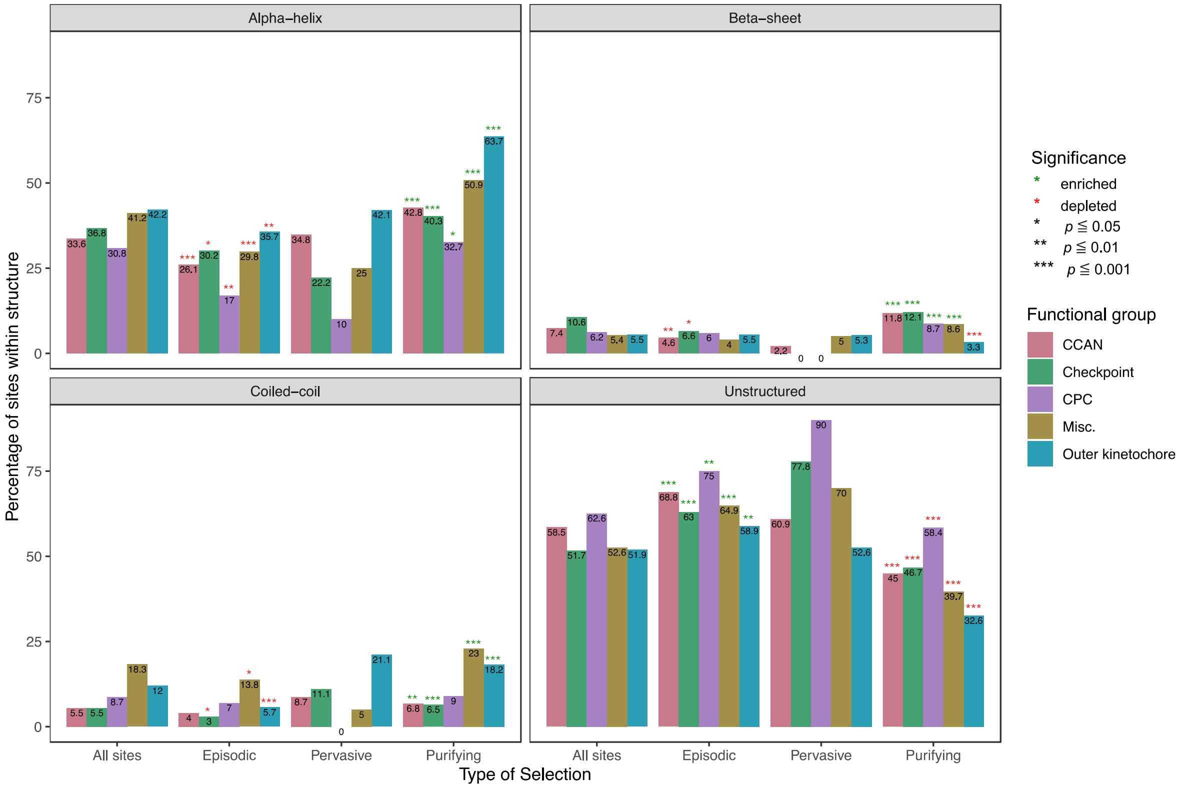


**Figure S9.** Percentage of sites within each secondary structure region broken down by functional grouping. The bars are grouped on the x-axis by type of selection (episodic diversifying, pervasive diversifying, or purifying) and the "all sites" group is the percentage of sites within the particular structure region regardless of selection. Bars are colored by functional group. Significant enrichment of selection type within a particular structure is denoted with a green asterisk; depletion is denoted with a red asterisk.

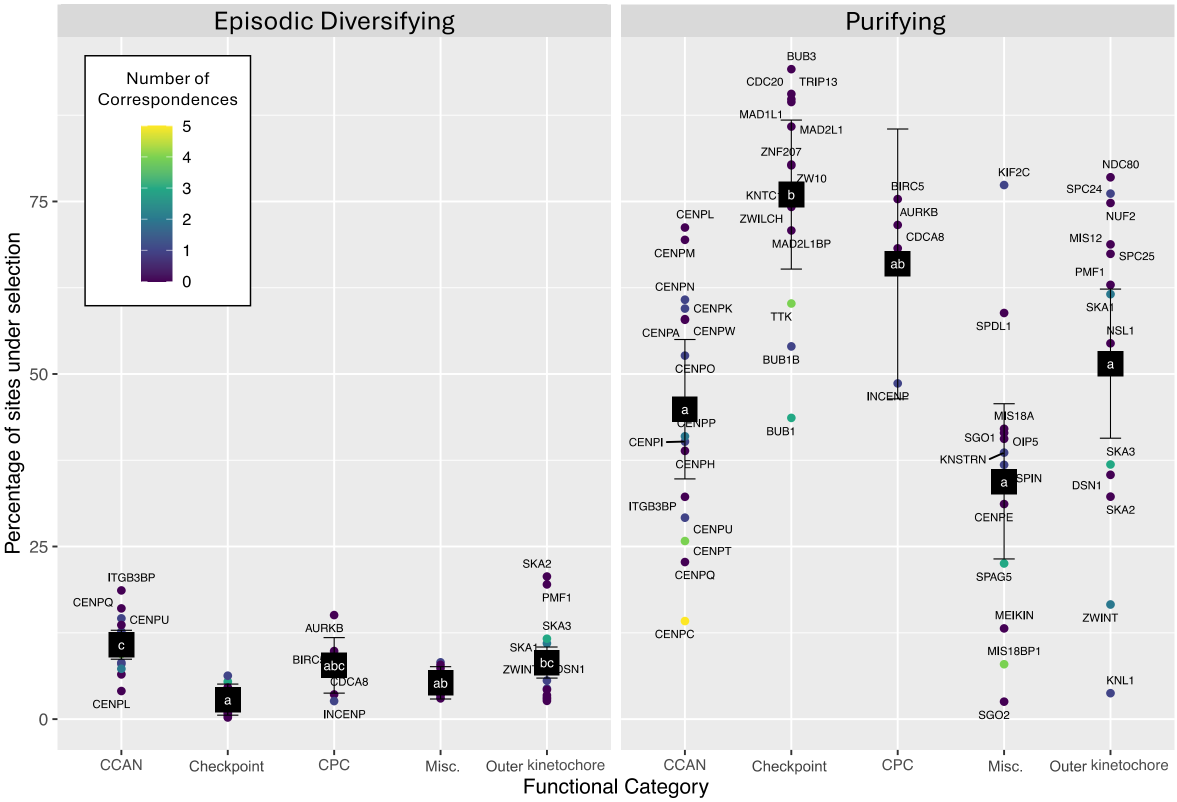


**Figure S10.** The percentage of sites under selection in each gene grouped according to functional category. Genes are color coded to represent the number of sites with corresponding evolutionary patterns with other genes in the study. Note that the data is from the same analysis as that depicted in figure 2. Black squares are the least squares mean for each functional category and bars are the 95% confidence interval around the means. Letters represent significance groups according to a pairwise Tukey test comparing the means of all five groups.

**Table S1.** RefSeq (O'Leary et al. 2016) accession numbers for the genomic protein sequences utilized in this study including 70 mammalian species and 4 outgroups.

| Species | RefSeq Accession | Species | RefSeq Accession |
|---|---|---|---|
| **Acinonyx jubatus** | GCF_3709585.1 | **Homo sapiens** | GCF_000001405.39 |
| **Ailuropoda melanoleuca** | GCF_2007445.1 | **Hylobates moloch** | GCF_9828535.2 |
| **Alligator mississippiensis** | GCF_281125.3 | **Ictidomys tridecemlineatus** | GCF_236235.1 |
| **Anolis carolinensis** | GCF_90745.1 | **Lepisosteus oculatus** | GCF_242695.1 |
| **Aotus nancymaae** | GCF_952055.2 | **Lontra canadensis** | GCF_10015895.1 |
| **Arvicanthis niloticus** | GCF_11762505.1 | **Loxodonta africana** | GCF_1905.1 |
| **Balaenoptera acutorostrata** | GCF_493695.1 | **Macaca mulatta** | GCF_3339765.1 |
| **Bison bison** | GCF_754665.1 | **Mandrillus leucophaeus** | GCF_951045.1 |
| **Bos taurus** | GCF_2263795.1 | **Manis javanica** | GCF_1685135.1 |
| **Callorhinus ursinus** | GCF_3265705.1 | **Marmota marmota** | GCF_1458135.1 |
| **Camelus bactrianus** | GCF_767855.1 | **Microcebus murinus** | GCF_165445.2 |
| **Canis lupus familiaris** | GCF_2285.3 | **Monodelphis domestica** | GCF_2295.2 |
| **Capra hircus** | GCF_1704415.1 | **Monodon monoceros** | GCF_5190385.1 |
| **Carlito syrichta** | GCF_164805.1 | **Mus musculus** | GCF_1635.27 |
| **Castor canadensis** | GCF_1984765.1 | **Mustela putorius** | GCF_215625.1 |
| **Cavia porcellus** | GCF_151735.1 | **Myotis lucifugus** | GCF_147115.1 |
| **Ceratotherium simum** | GCF_283155.1 | **Ochotona princeps** | GCF_292845.1 |
| **Chinchilla lanigera** | GCF_276665.1 | **Odocoileus virginianus** | GCF_2102435.1 |
| **Chlorocebus sabaeus** | GCF_409795.2 | **Orcinus orca** | GCF_331955.2 |
| **Chrysochloris asiatica** | GCF_296735.1 | **Ornithorhynchus anatinus** | GCF_4115215.1 |
| **Condylura cristata** | GCF_260355.1 | **Orycteropus afer** | GCF_298275.1 |
| **Cricetulus griseus** | GCF_223135.1 | **Oryctolagus cuniculus** | GCF_3625.3 |
| **Dasypus novemcinctus** | GCF_208655.1 | **Otolemur garnettii** | GCF_181295.1 |
| **Delphinapterus leucas** | GCF_2288925.2 | **Ovis aries** | GCF_2742125.1 |
| **Desmodus rotundus** | GCF_2940915.1 | **Panthera tigris** | GCF_464555.1 |
| **Echinops telfairi** | GCF_313985.2 | **Pan troglodytes** | GCF_2880755.1 |
| **Elephantulus edwardii** | GCF_299155.1 | **Phascolarctos cinereus** | GCF_2099425.1 |
| **Enhydra lutris** | GCF_2288905.1 | **Pongo abelii** | GCF_2880775.1 |
| **Eptesicus fuscus** | GCF_308155.1 | **Rattus norvegicus** | GCF_1895.5 |
| **Equus caballus** | GCF_2863925.1 | **Sarcophilus harrisii** | GCF_902635505.1 |
| **Eumetopias jubatus** | GCF_4028035.1 | **Suricata suricatta** | GCF_6229205.1 |
| **Felis catus** | GCF_181335.3 | **Sus scrofa** | GCF_3025.5 |
| **Fukomys damarensis** | GCF_12274545.1 | **Trichechus manatus** | GCF_243295.1 |
| **Gallus gallus** | GCF_2315.6 | **Tupaia chinensis** | GCF_334495.1 |
| **Globicephala melas** | GCF_6547405.1 | **Ursus arctos** | GCF_3584765.1 |
| **Gorilla gorilla** | GCF_8122165.1 | **Vicugna pacos** | GCF_164845.3 |
| **Heterocephalus glaber** | GCF_247695.1 | **Vombatus ursinus** | GCF_900497805.2 |

**Table S2.** Site-specific breakdown of corresponding sites. The columns labeled "Gene 1" and "Gene 2" contain the gene names in which the two corresponding sites reside. If the gene names are the same, the correspondence was internal. The human sequence site number and the site number corresponding to the overall gene alignment for each site are located in the second and sixth columns. The "Residue change" column contains information regarding the sequence changes in the species under selection. The amino acid before the arrow is the most common at that particular site and the amino acids after the arrow represent the residues present in the evolving species (last column). Highly diverse sites where a "most common" residue at that site was hard to identify are highlighted. Finally the secondary structure prediction at each site is reported, where NA represents no predicted structure.

| Gene 1 | Human site # (Alignment site #) | Residue change (human -> species AA) | SS pred | Gene 2 | Human site # (Alignment site #) | Residue change (human -> species AA) | SS pred | Correlating Species |
|---|---|---|---|---|---|---|---|---|
| **AURKB** | 6 (100) | E -> P, G, G, T, K | NA | **AURKB** | 7 (101) | L -> R, R, R, F, S | NA | G. gorilla, H. moloch, M. javanica, S. suricatta |
| **BUB1** | 507 (579) | V -> T, E, T | NA | **CENPC** | 59 (209) | V -> M, I, I | NA | C. syrichta, C. cristata, D. rotundus |
| **BUB1** | 154 (212) | A -> V, V, S, V | NA | **CENPP** | 211 (357) | A -> P, P, T, T | NA | C. asiatica, C. cristata, D. rotundus, T. chinensis |
| **BUB1** | 367 (427) | C -> Y, H, R | NA | **SPAG5/Astrin** | 163 (284) | P -> S, N, I | NA | A. jubatus, A. melanoleuca, O. garnettii |
| **BUB1B** | 509 (564) | S -> L, L, L | NA | **KNSTRN** | 295 (456) | L -> S, T, G | helix | C. syrichta, C. asiatica, C. griseus |
| **CENPC** | 927 (1128) | K -> R, S, R | sheet | **CENPP** | 23 (116) | E -> L, S, R | NA | C. porcellus, M. lucifugus, T. chinensis |
| **CENPC** | 599 (781) | G -> N, I, D | NA | **MIS18BP1** | 29 (29) | I -> V, V, V | NA | C. lanigera, D. rotundus, O. princeps |
| **CENPC** | 599 (781) | G -> N, I, D | NA | **MIS18BP1** | 631 (639) | Q -> H, S, H | NA | C. lanigera, D. rotundus, O. princeps |
| **CENPC** | 688 (882) | R -> T, T, T | NA | **ZWINT** | 247 (285) | Q -> E, H, W | NA | A. nancymaae, C. cristata, E. caballus |
| **CENPC** | 2 (117) | A -> C, L, C, L | NA | **CENPC** | 4 (120) | S -> A, F, G, E | NA | C. syrichta, M. murinus, O. afer, P. tigris |
| **CENPK** | 77 (198) | K -> Q, K, K | coiled coil | **SPC24** | 148 (197) | K -> R, H, R | NA | A. mississippiensis, A. carolinensis, O. afer |
| **CENPI** | 63 (153) | E -> G, G, R | helix | **TTK/Mps1** | 219 (258) | L -> R, Q, S | NA | M. javanica, O. cuniculus, O. garnettii |
| **CENPN** | 91 (154) | G -> R, E, K, E | NA | **CENPT** | 441 (520) | H -> I, K, V, V | NA | C. hircus, C. syrichta, C. asiatica, E. fuscus |
| **CENPO** | 296 (480) | M -> V, T, I, L | NA | **INCENP** | 845 (905) | T -> S, S, A, A | NA | C. lupus, C. syrichta, D. novemcinctus, D. rotundus |
| **CENPT** | 404 (483) | A -> S, S, T | NA | **SKA1** | 105 (192) | V -> D, E, A | NA | C. syrichta, C. asiatica, D. rotundus |
| **CENPT** | 328 (398) | D -> S, A, G | NA | **TTK/Mps1** | 365 (410) | L -> I, P, Q | NA | C. cristata, D. rotundus, M. lucifugus |
| **CENPT** | 452 (531) | P -> L, L, R | NA | **TTK/Mps1** | 365 (410) | L -> I, P, Q | NA | C. cristata, D. rotundus, M. lucifugus |
| **CENPU** | 38 (279) | C -> C, T, Y | NA | **HASPIN** | 762 (811) | C -> S, S, H | NA | C. cristata, D. novemcinctus, H. glaber |
| **KIF2C** | 145 (270) | V -> I, I, A | NA | **ZWINT** | 247 (285) | Q -> E, W, H | NA | A. nancymaae, E. caballus, C. cristata |
| **KNL1** | 2145 (2205) | D -> N, G, E | NA | **SKA3** | 197 (231) | L -> V, P, P | NA | A. melanoleuca, E. caballus, M. murinus |
| **KNTC1** | 2204 (2283) | F -> L, K, K, S | helix | **KNTC1** | 2208 (2295) | L -> L, S, C, Q | NA | A. nancymaae, C. sabaeus, E. edwardii, G. melas |
| **MIS18BP1** | 29 (29) | I -> V, V, V | NA | **MIS18BP1** | 631 (639) | Q -> H, S, H | NA | C. lanigera, |

| | | | | | | | | |
|---|---|---|---|---|---|---|---|---|
| | | | | | | | | D. rotundus,<br>O. princeps |
| **MIS18BP1** | 487 (491) | K -> R, E, E | NA | **SKA3** | 254 (292) | T -> P, R, K | NA | C. simum,<br>I. tridecemlineatus,<br>O. afer |
| **MIS18BP1** | 920 (943) | R -> K, N, K | helix | **SPAG5/Astrin** | 125 (245) | Y -> S, D, K | NA | E. caballus, O. afer,<br>S. suricatta |
| **SKA1** | 133 (225) | S -> I, A, F | NA | **SKA3** | 16 (48) | S -> V, S, V | helix | C. cristata,<br>M. javanica,<br>S. suricatta |
| **SPAG5/Astrin** | 653 (788) | T -> K, L, V | helix | **TTK/Mps1** | 330 (373) | V -> I, L, I | NA | O. garnettii,<br>T. manatus,<br>T. chinensis |
| **ZWINT** | 275 (317) | N -> R, A, S | NA | **ZWINT** | 268 (310) | L -> P, V, C | NA | M. mulatta,<br>M. javanica,<br>M. lucifugus |

**Table S3.** The human sequence included in each gene's alignment. For our analysis, one human sequence representing the most highly expressed transcript in human cells that generates the full length annotated protein and/or maximizes sequence length was chosen to be included in each gene's alignment. For each gene, the table includes the transcript ID according to Ensembl (Cunningham et al. 2022) and the corresponding RefSeq (O'Leary et al. 2016) accession number for the human transcript used.

| Gene | Transcript ID | RefSeq Accession | Gene | Transcript ID | RefSeq Accession |
|---|---|---|---|---|---|
| **AURKB** | ENST00000585124.5 | NM_004217.4 | **KNSTRN** | ENST00000249776.12 | NM_033286.4 |
| **BIRC5** | ENST00000350051.7 | NM_001168.3 | **KNTC1** | ENST00000333479.11 | NM_014708.6 |
| **BUB1** | ENST00000302759.10 | NM_004336.5 | **MAD1L1** | ENST00000265854.12 | NM_001013836.2 |
| **BUB1B** | ENST00000287598.10 | NM_001211.6 | **MAD2L1** | ENST00000333047.9 | NM_002358.4 |
| **BUB3** | ENST00000368858.9 | NM_001007793.3 | **MAD2L1BP** | ENST00000372171.4 | NM_014628.3 |
| **CDC20** | ENST00000310955.10 | NM_001255.3 | **MEIKIN** | ENST00000442687.6 | NM_001303622.2 |
| **CDCA8** | ENST00000373055.5 | NM_001256875.2 | **MIS12** | ENST00000381165.3 | NM_024039.3 |
| **CENPA** | ENST00000335756.8 | NM_001809.4 | **MIS18A** | ENST00000290130.3 | NM_018944.3 |
| **CENPC** | ENST00000273853.11 | NM_001812.4 | **MIS18BP1** | ENST00000310806.8 | NM_018353.5 |
| **CENPE** | ENST00000380026.7 | NM_001286734.2 | **NDC80** | ENST00000261597.8 | NM_006101.3 |
| **CENPH** | ENST00000283006.6 | NM_022909.4 | **NSL1** | ENST00000366977.7 | NM_015471.4 |
| **CENPI** | ENST00000372927.5 | NM_006733.3 | **NUF2** | ENST00000271452.7 | NM_145697.3 |
| **CENPK** | ENST00000396679.5 | NM_022145.5 | **OIP5** | ENST00000220514.7 | NM_007280.2 |
| **CENPL** | ENST00000367710.7 | NM_001171182.2 | **PMF1** | ENST00000368277.3 | NM_007221.4 |
| **CENPM** | ENST00000215980.9 | NM_024053.5 | **SGO1** | ENST00000442720.5 | NM_001199255.3 |
| **CENPN** | ENST00000305850.10 | NM_001100624.3 | **SGO2** | ENST00000357799.8 | NM_152524.6 |
| **CENPO** | ENST00000380834.7 | NM_001322101.2 | **SKA1** | ENST00000285116.7 | NM_145060.4 |
| **CENPP** | ENST00000618653.1 | NM_001012267.3 | **SKA2** | ENST00000330137.12 | NM_182620.4 |
| **CENPQ** | ENST00000335783.3 | NM_018132.4 | **SKA3** | ENST00000314759.5 | NM_145061.6 |
| **CENPT** | ENST00000562787.6 | NM_025082.4 | **SPAG5/Astrin** | ENST00000321765.9 | NM_006461.4 |
| **CENPU** | ENST00000281453.9 | NM_024629.4 | **SPC24** | ENST00000592540.5 | NM_182513.4 |
| **CENPW** | ENST00000368328.4 | NM_001012507.4 | **SPC25** | ENST00000282074.6 | NM_020675.4 |
| **DSN1** | ENST00000373750.8 | NM_001145315.2 | **SPDL1** | ENST00000265295.8 | NM_017785.5 |
| **HASPIN** | ENST00000325418.5 | NM_031965.2 | **TRIP13** | ENST00000166345.7 | NM_004237.4 |
| **INCENP** | ENST00000394818.7 | NM_001040694.2 | **TTK/Mps1** | ENST00000369798.6 | NM_003318.5 |
| **ITGB3BP** | ENST00000271002.14 | NM_014288.5 | **ZNF207** | ENST00000394673.6 | NM_001032293.3 |
| **KIF2C** | ENST00000372224.8 | NM_006845.4 | **ZW10** | ENST00000200135.7 | NM_004724.4 |
| **KNL1** | ENST00000399668.7 | NM_144508.5 | | | |
